# Whole-Brain MRSI Metabolic Connectomics Reveals Network Disruptions and Altered Gradient Coherence in Psychosis Risk

**DOI:** 10.64898/2026.11.09.765372

**Authors:** Federico Lucchetti, Edgar Céléreau, Raoul Jenni, Teya Petrova, Lilith Abrahamyan, Philippe Conus, Patric Hagmann, Yasser Alemán-Gómez, Antoine Klauser, Paul Klauser

## Abstract

Brain network alterations preceding first-episode psychosis often escape connectomic approaches based on conventional neuroimaging, such as diffusion or functional magnetic resonance imaging (MRI). Whole-brain magnetic resonance spectroscopic imaging (MRSI) offers a potentially more sensitive alternative: by mapping neurometabolites across the brain, it enables the construction of metabolic connectomes that may capture very early and subtle disruptions of brain network properties. We combined whole-brain proton MRSI, diffusion MRI, and resting-state functional MRI in a sample of 29 individuals meeting DSM-5 criteria for Attenuated Psychosis Syndrome (APS) or Schizotypal Personality Disorder (SCZT), pooled into a single High Risk for Psychosis (HR) group, and 30 healthy controls (HC) to construct metabolic, structural, and functional connectomes. Network-based statistics identified a significant metabolic connectivity subnetwork with higher connectivity in HR than in HC, whereas no corresponding structural or functional connectivity differences were detected. Nodes of this subnetwork had modestly higher structural strength but were not consistently enriched among structural hubs, and no significant regional abnormalities in individual metabolite concentrations were detected. Eigenmode analysis of metabolic connectivity revealed a low-dimensional organization spanning a posterior unimodal-to-transmodal axis and a transmodal-to-limbic axis. Path analysis revealed higher bilateral energy in HR, indicating reduced coherence along a metabolic gradient that parallels the neurodevelopmental sensorimotor- to-association axis. These findings identify whole-brain metabolic network organization derived from MRSI as a systems-level phenotype of psychosis risk and raise the possibility that it captures altered development of large-scale cortical organization.

## Introduction

Schizophrenia is increasingly understood as a disorder of distributed brain network organization, rather than a pathology of isolated regions. Neuroimaging studies have consistently reported widespread abnormalities in the structural and functional network of the brain Kelly et al. (2018); Klauser et al. (2017a); Dong et al. (2018); van den Heuvel et al. (2013). However, by the time schizophrenia is clinically established, substantial neurobiological changes may already have occurred, limiting opportunities for early intervention. Indeed, psychosis spectrum disorders usually emerge in late adolescence or early adulthood and may progress in a psychosis continuum from a high risk (HR) state to a first-episode psychosis and, in some cases, to an established psychotic disorder such as chronic schizophrenia Lieberman et al. (2019). A primary objective is to identify HR individuals, with the aim of offering them an early intervention that targets the mechanisms of the disorder. In this context, the identification of brain biomarkers for HR individuals plays a central role. Nevertheless, no imaging modality has yet been able to identify clinically significant brain alterations that would aid in early detection or guide the development of novel interventions Kraguljac et al. (2021); Fuentes-Claramonte et al. (2024). The present study attempts to addresses this need by characterizing brain metabolic network organization in HR, as a step toward identifying individuals most likely to deteriorate, and refining early intervention strategies before the onset of a first psychotic episode.

Structural and functional magnetic resonance imaging (MRI) connectomics have transformed our understanding of large-scale brain organization by revealing principles such as modular architecture, hub organization, and rich-club structure Sporns and Kötter (2004); Van Den Heuvel and Sporns (2011). Diffusion MRI (dMRI) provides estimates of structural connectivity based on putative white-matter pathways, whereas resting-state functional (fMRI) captures statistical dependencies between regional hemodynamic signals. These approaches have been particularly influential in neuropsychiatry, where highly central hub regions are thought to be vulnerable to pathological perturbations because of their integrative role and high metabolic demands Bullmore and Sporns (2012); van den Heuvel and Sporns (2013); Crossley et al. (2014); Van Den Heuvel and Sporns (2019); Lebel et al. (2008). However, both structural and functional connectivity remain indirect with respect to the biochemical processes that sustain neural signaling Logothetis (2008); Jones et al. (2013), leaving the metabolic substrate of brain network organization largely unresolved.

Molecular and metabolic imaging approaches offer a more biology-proximal perspective. Positron emission tomography (PET) uses radiotracers to image specific molecular targets and biological processes, typically one per tracer Phelps (2000); Hansen et al. (2022), whereas proton magnetic resonance spectroscopic imaging (MRSI) measures multiple endogenous brain metabolites simultaneously, without injected tracers Öz et al. (2014); Maudsley et al. (2021). Recent advances in fast whole-brain 3D proton MRSI now make it possible to map multiple metabolites non-invasively across the entire brain within clinically feasible scan times Bogner et al. (2021); Klauser et al. (2022, 2024), and to embed this information into a network framework Lucchetti et al. (2025); Liu et al. (2026). By characterizing large-scale patterns of biochemical similarity across brain regions, metabolic connectomics provides a complementary framework for investigating not only where pathology manifests within brain networks, but also whether vulnerable subnetworks are embedded within altered neurometabolic organization.

Recent work using whole-brain MRSI has shown that within-subject metabolic connectivity (MC), constructed from regional profiles of major brain metabolites, captures a coherent large-scale axis of biochemical organization Lucchetti et al. (2025). At 3T MRI field strengths, these profiles are primarily defined by total N-acetylaspartate plus N-acetylaspartylglutamate (tNAA), total creatine (tCr), glutamate plus glutamine (Glx), choline-containing compounds (Cho), and myo-inositol (Ins). While these signals are not direct readouts of brain bioenergetic dynamics, they are embedded in energetically demanding pathways involving neurotransmitter cycling, glial and neuronal metabolism, membrane turnover, and phosphocreatine buffering Harris et al. (2012); Magistretti and Allaman (2015). MC has further been shown to capture biologically meaningful aspects of tissue composition and large-scale brain organization, as evidenced by its close alignment with cytoarchitectonic similarity and gene co-expression in healthy individuals Lucchetti et al. (2025). Metabolic connectivity also exhibits a coherent dominant gradient of brain organization, linking caudal unimodal territories with transmodal and limbic regions. A similar principal gradient has been described in functional connectivity Margulies et al. (2016) and shown to be altered across several neuropsychiatric disorders Park et al. (2022), consistent with its proposed role in supporting cognitive flexibility Nashed et al. (2025). This axis closely resembles the sensorimotor- to-association (S-A) axis described across structural, functional, and developmental neuroimaging, along which cortical myelination and maturation unfold during typical development Sydnor et al. (2021); Whitaker et al. (2016); Paquola et al. (2019b,a). Recent multispecies evidence further indicates that the S-A axis is conserved across primates Buckner and Krienen (2013), emerges through spatially organized and competing molecular developmental programs Tsyporin et al. (2026), reinforcing its developmental basis. Together, these observations suggest that large-scale brain gradients may retain a spatial signature of coordinated neurodevelopment. Just as the concentric growth rings of an expanding tree provide a spatial record of its development and can reveal periods of disruption, the coherence of spatial progression along large-scale brain gradients may similarly reflect the integrity of brain maturation. Here, we acquired whole-brain MRSI, diffusion MRI, and resting-state fMRI in healthy controls and HR participants, and constructed metabolic, structural, and functional connectivity matrices for each subject. We tested three hypotheses: first, that HR would show alterations in metabolic connectivity even in the absence of detectable structural or functional connectivity differences; second, that these metabolic alterations would preferentially involve subnetworks converging on structurally central hubs; and third, that HR would show reduced coherence of the large-scale metabolic connectivity gradient, reflecting a less orderly spatial organization along a putative neurodevelopmental axis.

## Results

### Connectome construction

We first constructed subject-level structural, functional, and metabolic connectomes in matched subject-specific volumetric parcellations (Fig. 1). MRI and MRSI acquisition protocols are detailed in MRI image acquisition, and anatomical preprocessing, Chimera parcellation, and modality-specific connectome construction are described in Preprocessing and connectome construction. Briefly, the primary analyses used the LFMIHIFS-3 parcellation, which combines the Lausanne scale 3 cortical parcellation with subcortical, cerebellar, and brainstem subdivisions. Two additional Chimera schemes, based on MIST-197 and Schaefer-200 cortical subdivisions with the same subcortical definitions, were used as parcellation controls and assess the influence of the parcellation scheme on the results that follow.

**Figure 1:**
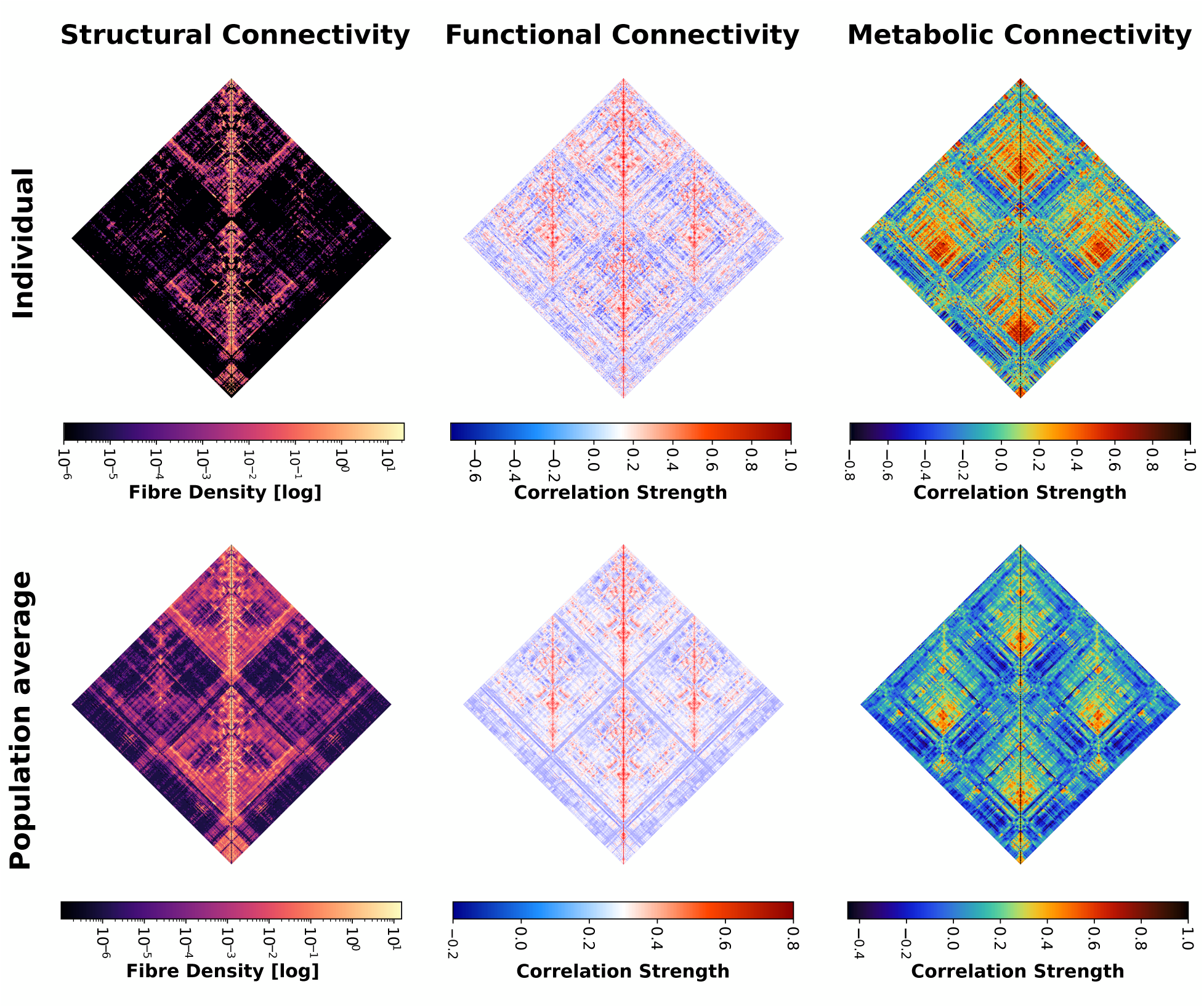
Representative individual (top row) and population-average (bottom row) matrices are shown for structural connectivity, resting-state BOLD-derived functional connectivity, and MRSI-derived metabolic connectivity. All modalities were mapped to the same subject-specific Chimera LFMIHIFS-3 parcellation.

For each participant, diffusion MRI yielded structural connectivity matrices (SC), resting-state fMRI yielded functional connectivity matrices (FC), and whole-brain ^1^H-MRSI yielded metabolic connectivity matrices (MC). We use the term *metabolic connectivity* to denote an edge-wise relationship between brain regions obtained by comparing their representations over the metabolic signal domain encoded in the chemical-shift dimension measured by MRSI. This parallels functional connectivity, in which regional blood-oxygen-level-dependent (BOLD) signals are compared over the temporal domain, and structural connectivity, in which anatomical connection evidence is inferred from diffusion information distributed along physical white-matter trajectories.

Because MRSI data were acquired with two sequence implementations on the same MRI platform, namely the original 3D ^1^H-FID-MRSI sequence and the ECCENTRIC 3D ^1^H-FID-MRSI sequence, MC matrices were harmonized across sequences using neuroCombat while preserving variance associated with age, sex, and diagnostic group (Cross-sequence harmonization; Supplementary Material REF).

After modality-specific quality control, the retained samples comprised 30 healthy controls (HC) and 27 individuals at High Risk for Psychosis (HR) for metabolic connectivity, 31 HC and 29 HR participants for structural connectivity, and 27 HC and 26 HR participants for functional connectivity.

Representative individual-level and population-average matrices are shown in Figure 1 for the LFMIHIFS-3 parcellation. These matrices provided the basis for subsequent group comparisons between healthy controls and HR participants across modalities in the following sections.

### Metabolic connectivity alterations

We tested whether HR participants and healthy controls (HC) differed in their subject-level matrices using the network-based statistic (Network-based analysis). In the LFMIHIFIS-3 parcellation, the model adjusted for sex and age identified an HR *>* HC component at the reporting threshold (*t* = 3.25) comprising 68 nodes and 120 edges, which was significant after correction for the two tested directions (two-direction-corrected *p* = 0.0260; uncorrected directional *p* = 0.0130; Fig. 2**A**). Because MRSI acquisition sequence had already been harmonized with neuroCombat, we additionally included sequence as a nuisance covariate as a sensitivity analysis. This model yielded the same direction and a component of similar size (66 nodes and 109 edges; corrected *p* = 0.0272; uncorrected directional *p* = 0.0136), which was present across all tested primary thresholds (*t* = 3.00–4.00), indicating that the LFMIHIFIS-3 result was not dependent on residual sequence effects.

**Figure 2:**
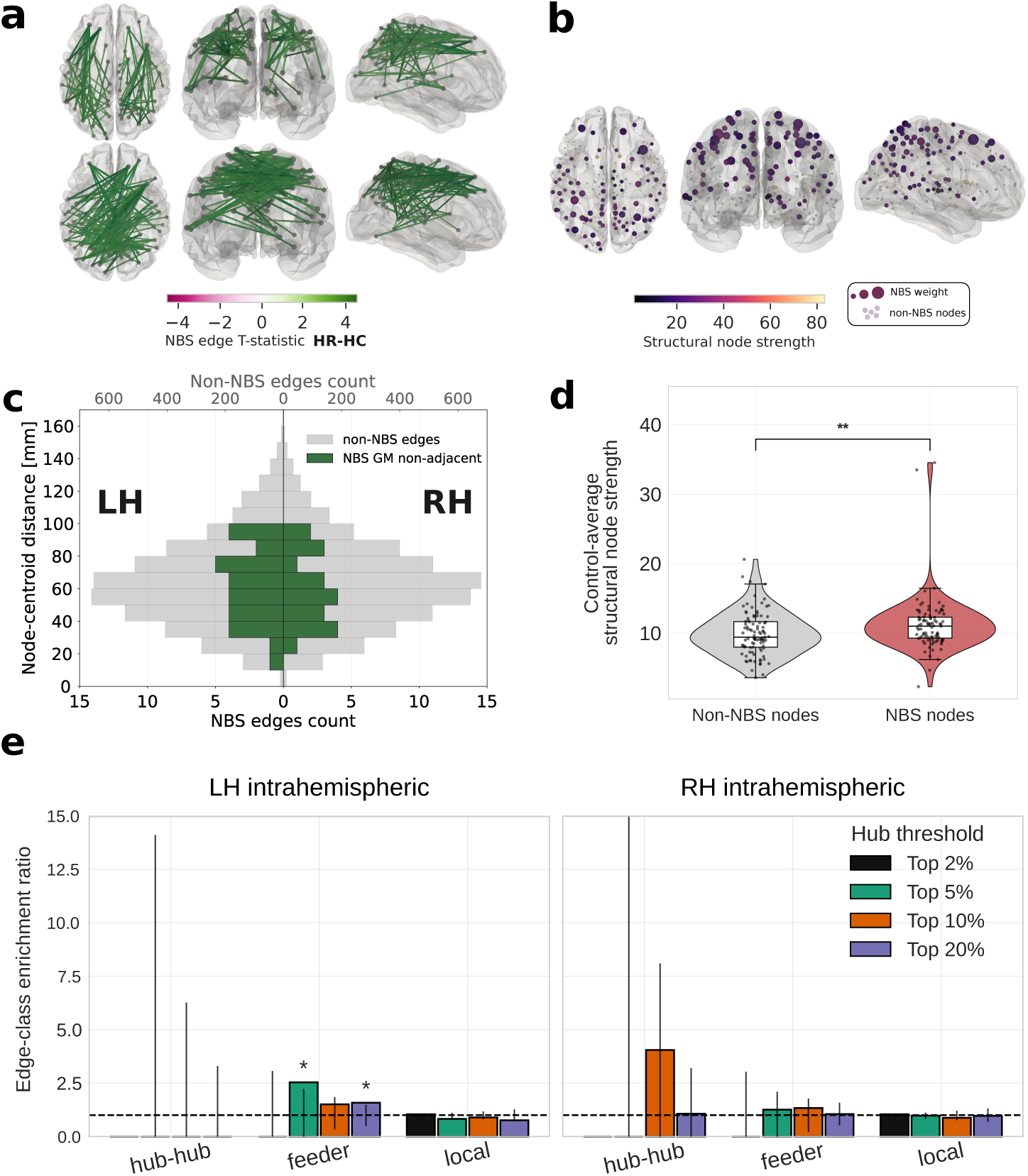
Structural context of the metabolic NBS subnetwork. textbfA, Significant metabolic NBS subnetwork for the HR *>* HC MC contrast (primary NBS threshold *t* = 3.25; sex- and age-adjusted, two-direction-corrected *p* = 0.0260), with intrahemispheric NBS edges shown at the top and interhemispheric NBS edges at the bottom. **B**, Control-average structural node strength projected onto the same parcellation, with nodes incident to at least one NBS edge identified in panel A highlighted and scaled by their NBS load. **C**, Distribution of intrahemispheric edge lengths (Euclidean distance between parcel centroids) for NBS edges linking non-adjacent grey-matter parcels (green; bottom axis) and for all non-NBS edges (grey; top axis), shown separately for the left (LH) and right (RH) hemispheres. The two count axes are scaled differently for visibility; interhemispheric NBS edges (*n* = 66) are not shown. **D**, Structural node strength in NBS and non-NBS nodes. **E**, Enrichment of NBS edges by structural edge class across hub definitions, shown separately for intrahemispheric edges of the left (LH) and right (RH) hemispheres, with hubs defined within each hemisphere. Hub–hub, feeder, and local edges connect two hubs, one hub and one non-hub, or two non-hubs, respectively. Bars show enrichment relative to random intrahemispheric edge sets of equal size; intervals indicate the 95% null range and stars denote empirical significance.

At *t* = 3.25, the sequence-adjusted MFMIHIFIS-197 analysis showed an HR *>* HC component that did not cross the corrected significance threshold (37 nodes, 47 edges; *p* = 0.0544), whereas the SFMIHIFIS-200 component was significant (77 nodes, 122 edges; *p* = 0.0280). Without MRSI-sequence adjustment, the corresponding p-values were 0.0456 for MFMIHIFIS-197 (41 nodes, 54 edges) and 0.0248 for SFMIHIFIS-200 (79 nodes, 136 edges), again in the HR *>* HC direction. This effect was specific to MC. We applied the same covariate-adjusted NBS framework to the SC and FC matrices. For SC, 5,000-permutation analyses across primary thresholds *t* = 3.00, 3.25, 3.50, 3.75, and 4.00 detected no significant component in either direction for any parcellation. The smallest two-direction-corrected component-level p-values were 0.7544 for LFMIHIFIS-3, 0.7980 for MFMIHIFIS-197, and 0.3500 for SFMIHIFIS-200. The corresponding FC analyses likewise detected no significant component in either direction; the smallest corrected p-values were 0.1140, 0.2392, and 0.0924, respectively. Thus, within the present sample, group differences were detectable at the level of MC but not in the corresponding SC and FC. Results for all modalities, parcellations, adjustment models, and primary thresholds are summarized in Supplementary Table S1.

### Metabolic NBS nodes show no consistent convergence on structural hubs

We then tested our second hypothesis, that the altered metabolic subnetwork would preferentially involve structural hubs (Structural context of the metabolic NBS component). Nodes incident to edges in the significant MC NBS component were projected onto the control-average structural connectome, and structural node strength was computed for each parcel (Fig. 2**B**). NBS-involved nodes had higher structural strength than non-NBS nodes (two-sided Mann–Whitney *p* = 0.0008; rank-biserial correlation = 0.296; Fig. 2**D**). NBS node load also correlated positively with structural strength (Spearman *ρ* = 0.235, parametric *p* = 0.0012, permutation *p* = 0.0018), indicating a modest association between metabolic alteration and structural strength.

We next tested whether NBS-involved nodes were enriched among strength-defined structural hubs, analysing each hemisphere separately with hubs defined within that hemisphere. No hub-node enrichment was detected at any hub definition in either hemisphere (all empirical *p* ≥ 0.16). At the edge level, intrahemispheric NBS edges were classified as hub–hub, feeder, or local (Fig. 2**E**). In the left hemisphere, feeder edges were enriched at the 5% (8 observed; enrichment ratio = 2.53; empirical *p* = 0.011), 10% (12 observed; enrichment ratio = 2.01; *p* = 0.009), and 20% definitions (19 observed; enrichment ratio = 1.88; *p* = 0.001). In the right hemisphere, no edge class was enriched at any definition (all *p* ≥ 0.078). No hub–hub enrichment was detected in either hemisphere; the right hemisphere contained a single hub–hub NBS edge at the 20% definition. Local edges were not enriched at any definition. Across the three parcellations and five primary thresholds, hub-node and hub–hub enrichment remained largely absent, whereas feeder enrichment recurred but switched hemisphere between parcellations (Supplementary Tables S2 and S3). Thus, the second hypothesis was not supported: hub-node and hub–hub enrichment were absent, and the edge-level effect was a feeder-edge preference whose lateralization was not consistent across parcellations.

We next examined the spatial scale of the altered connections (Structural context of the metabolic NBS component). The NBS edges spanned the whole network, from short- to medium-range parcel pairs (Fig. 2**C**), and 96% of them linked parcels that were not adjacent in grey matter. Within hemispheres, most NBS edges (63%) linked parcels separated by two to four adjacency steps along grey matter (median three). Because MC decreased with inter-parcel distance within each hemisphere in controls (Spearman *ρ* = −0.40 and −0.35 in the left and right hemispheres, respectively), these edges mostly connected regions that are normally weakly similar, but were more similar in HR.

We additionally tested whether the metabolic connectivity alteration was accompanied by abnormal local metabolism. This test was motivated by evidence that hubs form a densely interconnected cortical core and are disproportionately implicated across brain disorders (Hagmann et al., 2008; Crossley et al., 2014), together with evidence for marked regional variation in aerobic glycolysis (Vaishnavi et al., 2010) and prior reports of metabolite abnormalities in psychosis risk (Wang et al., 2020; Romeo et al., 2020). To assess this possibility, we performed parcel-wise comparisons of individual metabolite levels between HR participants and healthy controls (Parcel-wise metabolite analysis). Parcel-wise metabolite profiles were first harmonized across MRSI acquisition sequences using neuroCombat. Statistical testing was performed independently for each metabolite and corrected for multiple comparisons using the Benjamini–Hochberg false discovery rate procedure. No parcels survived correction for any metabolite (all FDR-adjusted *q* ≥ 0.586). These findings suggest that the observed group differences are not driven by focal abnormalities in absolute metabolite concentrations, but rather emerge at the level of inter-regional metabolic organization.

The same covariate-adjusted group-comparison framework was repeated using the additional Chimera parcellation schemes based on MIST-197 and Schaefer-200 cortical subdivisions. These complementary analyses identified effects in the same HR *>* HC direction, with sensitivity to parcellation and explicit MRSI-sequence adjustment reported in the Supplementary Material.

### Metabolic connectivity modes and gradients

To represent high-dimensional MC organization in a low-dimensional coordinate system, we applied diffusion embedding to the control-average MC matrix (Gradient analysis). The first two embedding dimensions, referred to as metabolic connectivity (MC) modes *g*_1_ and *g*_2_, revealed interpretable spatial axes: *g*_1_ varied predominantly along a rostro-caudal axis, whereas *g*_2_ captured a medial-lateral contrast (Fig. 3**a,b**).

**Figure 3:**
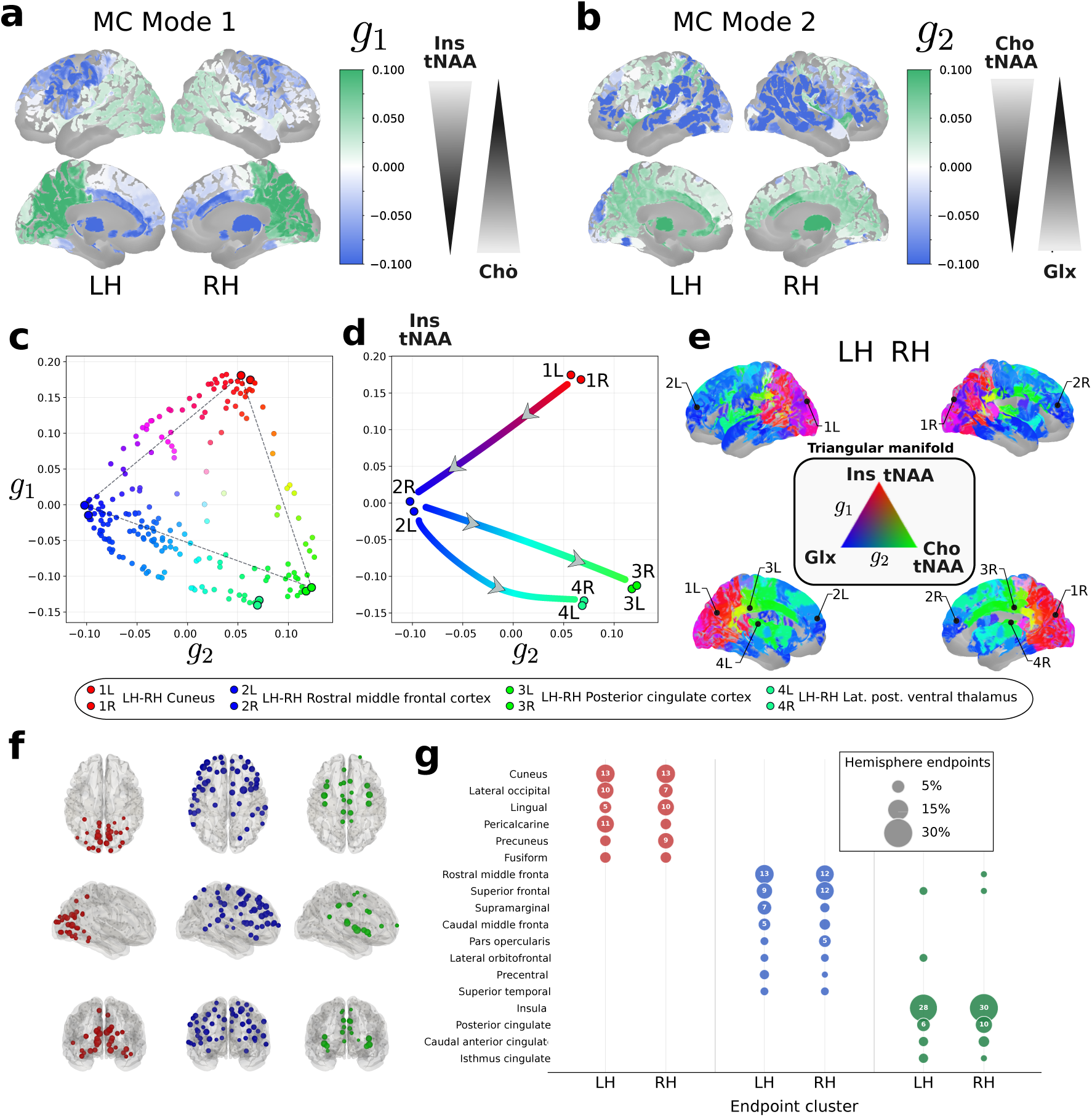
Metabolic connectivity modes define an intrinsic metabolic coordinate system. **a**, First metabolic connectivity (MC) mode (*g*_1_) derived from diffusion embedding of the control-average MC matrix, shown in lateral and medial views of both hemispheres. **b**, Second MC mode (*g*_2_). The metabolites whose relative contrast most strongly drives each mode (the Ins/tNAA–Cho antagonism for *g*_1_, the Cho/tNAA–Glx antagonism for *g*_2_) are indicated next to the colour scale. **c**, Control-average parcel coordinates in MC-mode space and the fitted triangular manifold. **d**, Directed principal axes linking the representative cuneus, rostral middle frontal, posterior cingulate, and lateral posterior ventral thalamic endpoints. **e**, Barycentric triangular-manifold colors projected onto the cortical surface, with the representative group-level endpoints marked. **f**, Pooled anatomical locations of the three cortical endpoint classes obtained by fitting the triangular manifold independently for each subject–session and hemisphere. **g**, Regional composition of the subject-level posterior unimodal, association, and limbic/insular endpoint classes, with paired left- and right-hemisphere columns. Circle size indicates the proportion of hemisphere-specific endpoints assigned to each region.

To identify the metabolites underlying these modes, we correlated each mode with pairwise metabolite contrast maps in controls (Metabolite contrasts underlying the MC modes; Supplementary Table S4 and Table S5). *g*_1_ was dominated by an Ins/tNAA–Cho antagonism (Ins and tNAA relatively elevated, Cho relatively reduced, toward higher *g*_1_; Fig. 3**a**), whereas *g*_2_ was dominated by a Cho/tNAA–Glx antagonism (Cho and tNAA relatively elevated, Glx relatively reduced, toward higher *g*_2_; Fig. 3**b**). Common maps, which track shared metabolite level rather than relative contrast, were only weakly associated with either mode (all |*r*| ≤ 0.43), so both modes primarily reflect relative differences between metabolites.

In the two-dimensional MC embedding space, parcels occupied a triangular manifold (Fig. 3**c**). Its fitted vertices defined representative endpoint classes, and the directed contrasts between vertices defined the principal axes through MC emebdding space (Fig. 3**d**). Each parcel was assigned three barycentric coordinates expressing its relative proximity to the vertices. Encoding these coordinates as an RGB mixture provided a continuous triangular-manifold representation that can be projected back onto the cortical surface (Fig. 3**e**).

At the group level, the dominant red-to-blue axis linked posterior unimodal cortex, including bilateral cuneus, to transmodal association cortex, including bilateral rostral middle frontal regions. This axis runs predominantly along *g*_1_ and reflects the Ins/tNAA–Cho antagonism. A secondary axis extended from transmodal cortex toward limbic regions, encompassing posterior cingulate cortex. This axis runs predominantly along *g*_2_ and reflects the Cho/tNAA–Glx antagonism. A closely parallel branch could also be delineated toward the lateral posterior ventral thalamus (Fig. 3**d,e**). This organization recapitulates the unimodal-to-transmodal axis described in functional connectivity gradients, but here emerged from pairwise metabolic connectivity estimated from regional MRSI-derived metabolite profiles.

The endpoint locations shown in Fig. 3**f,g** were obtained by fitting the triangular manifold independently for each subject and hemisphere. At the individual level, however, the limbic- and thalamic-directed branches could not always be reliably distinguished. The thalamic branch was therefore excluded from the subject-level endpoint analysis, which focused on the three reproducible cortical endpoint classes. Despite inter-individual variability, these converged on a consistent set of territories. Posterior unimodal endpoints were concentrated in cuneus, lateral occipital, lingual, and pericalcarine cortex, with precuneus as the main non-sensory exception. Association end-points occurred most frequently in rostral middle frontal and superior frontal cortex, followed by supramarginal, caudal middle frontal, and pars-opercularis parcels. Limbic endpoints were predominantly located in insular cortex, together with posterior, anterior, and isthmus cingulate regions. The pooled anatomical distributions are shown in Fig. 3**f**, and their left- and right-hemisphere composition in Fig. 3**g**.

To visualize whether this organization was preserved in HR, the HR-average MC embedding was aligned to the control-derived triangular coordinate system (Fig. 4**a**). The resulting parcel values were displayed using a control-derived circular ordering, with controls shown as circles and HR participants as squares (Fig. 4**b**). This representation makes local differences in MC-mode position visible, but it is quantitatively misleading if interpreted as a unique sequential ordering of the parcels.

**Figure 4:**
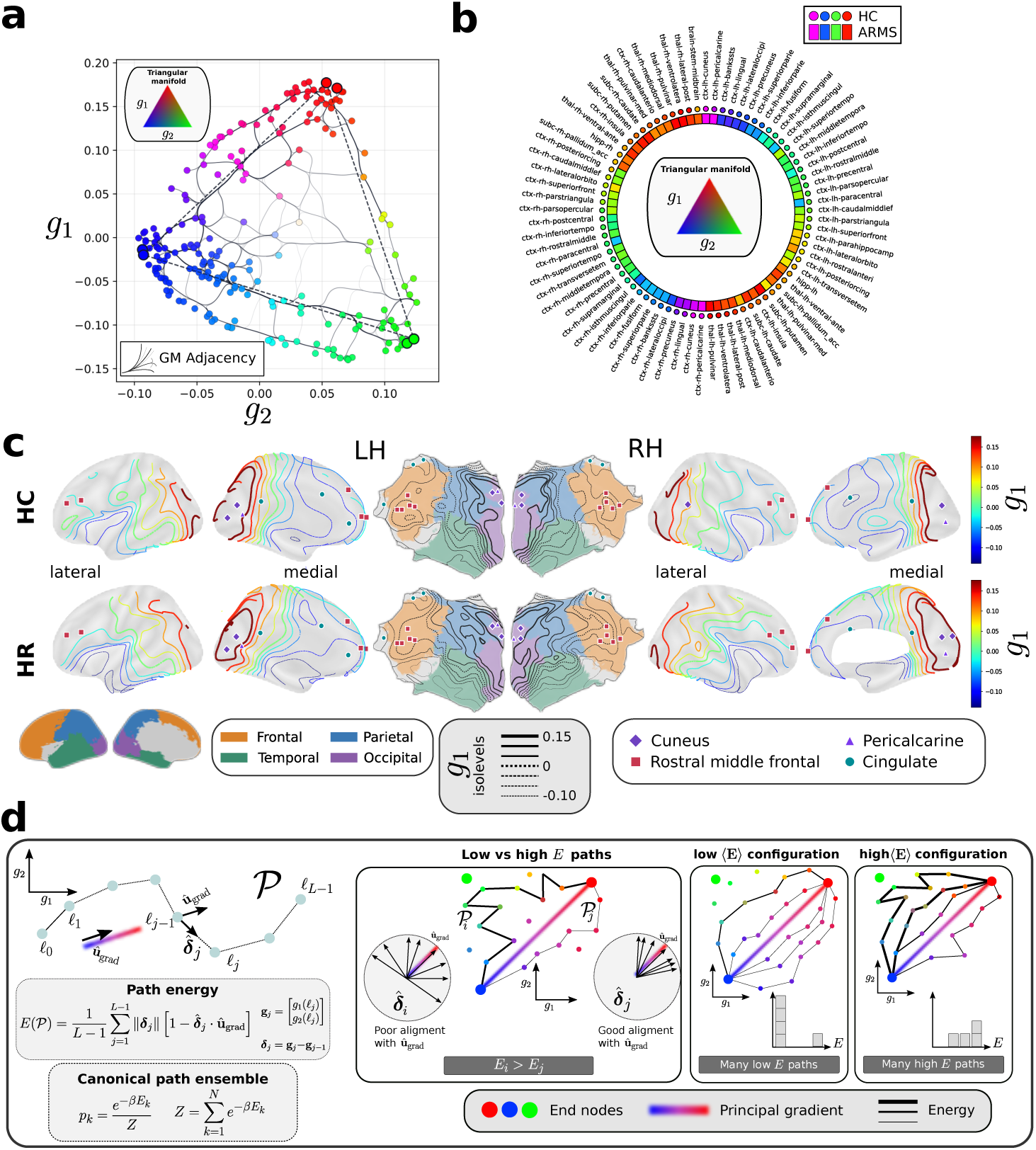
From visual MC-mode ordering to a grey-matter-constrained path ensemble. **a**, Control-average two-dimensional metabolic connectivity (MC) embedding with grey-matter adjacency overlaid; colors indicate position within the triangular manifold. **b**, Control-derived circular parcel ordering, with healthy controls (HC; circles) and HR participants (squares) colored by triangular-manifold coordinate. **c**, MC-mode *g*_1_ isolevels in HC (top) and HR (bottom), shown on lateral and medial cortical surfaces and on flattened left- and right-hemisphere maps. Contour colors encode *g*_1_ values and flatmap backgrounds identify cortical lobes. Markers identify cuneus, pericalcarine, rostral middle frontal, and cingulate parcels. **d**, Discrete path formulation showing segment-normalized path energy, normalized path weighting, and representative low- and high-energy path configurations.

The following point illustrates this limitation. The MC modes (*g*_1_*, g*_2_) define scalar fields over the cortical surface, whose isolevels appear as one-dimensional contours ((*g*_1_) shown on Fig. 4**c**). These contours are shown both on the folded three-dimensional cortical surface and, centrally, on flattened cortical maps to make their spatial progression across cortical territories more apparent. Although the isolevels define an ordered progression from low to high MC-mode values, they do not impose a unique ordering of the grey-matter parcels embedded within the cortical surface. Multiple distinct grey-matter-continuous sequences can therefore connect the same marked endpoint classes while progressing through the same MC-mode landscape. Each such path defines one possible anatomical ordering of parcels along the MC spatial gradient. This degeneracy of admissible orderings cannot be captured by the single sequence shown in Fig. 4**b**, motivating a framework that considers both the full set of grey-matter-continuous paths through the metabolic landscape and the coherence of the underlying gradient organization. The latter is already visually apparent in the flattened cortical maps of Fig. 4**c**, where successive isolevels appear more regular in controls and less coherent in HR.

### Path-based energy analysis

We addressed this by translating the continuous surface-gradient representation into a discrete, grey-matter-constrained path framework (Fig. 4**d**). The energy formulation was adapted from previous work showing that MC networks support smooth, monotonic rostro-caudal paths through MC gradient space and occupy an intermediate regime between geometric and random network organization (Lucchetti et al., 2025). Here, this formulation provides a means of quantifying how coherently anatomically admissible paths progress through the MC-mode landscape.

Low-energy paths were defined according to three complementary properties. First, they connect triangle-derived endpoint regions occupying distinct positions in the MC manifold. Second, they favor short successive displacements between grey-matter-adjacent parcels in MC-mode space. Third, these displacements remain aligned with the corresponding endpoint-defined direction. Path energy therefore penalizes both large local displacements and deviations from the principal MC-mode direction. A single subject-level value was obtained as the Boltzmann-weighted mean of the segment-normalized energies of all retained paths, with *β* = 1. Lower values indicate a smoother and more directionally coherent organization of the metabolic gradient. The path construction, energy function, and weighting are defined mathematically in Methods (Path-ensemble energy).

Segment-normalized path energy was lower in controls than in HR participants for both cortical trajectories (Fig. 5). Group-label permutation testing and spatial-null inference are detailed in Statistical analysis. After adjustment for sex and age, median energy was 0.0445 in controls and 0.0602 in HR for the left hemisphere (*p* = 0.0016) and 0.0458 and 0.0618, respectively, for the right hemisphere (*p* = 0.0036); both hemispheric tests remained significant after Benjamini–Hochberg correction for the two tested hemispheres (*q* = 0.0032 and *q* = 0.0036, respectively). Both effects were further evaluated against the Moran spatial-autocorrelation null and the adjacent-endpoint null (Statistical analysis), which respectively test whether the group difference could be explained by the intrinsic spatial autocorrelation of the metabolic gradients or by the precise anatomical placement of the selected endpoints. For the left hemisphere, the effect remained supported under both the Moran null (*p*_SA_,_moran_ = 0.0100) and the adjacent-endpoint null (*p*_SA_,_endpoint_ = 0.0247), yielding a conservative combined probability of *p*_SA_ = 0.0247. For the right hemisphere, the effect remained supported under the Moran null (*p*_SA_,_moran_ = 0.0120) but only at trend level under the adjacent-endpoint null (*p*_SA_,_endpoint_ = 0.0353), yielding a combined probability of *p*_SA_ = 0.0353. These findings indicate that HR-related MC alterations extend beyond localized edge differences and are associated with less directionally coherent progression through the cortical MC-mode landscape, with somewhat stronger null-model robustness in the left hemisphere.

**Figure 5:**
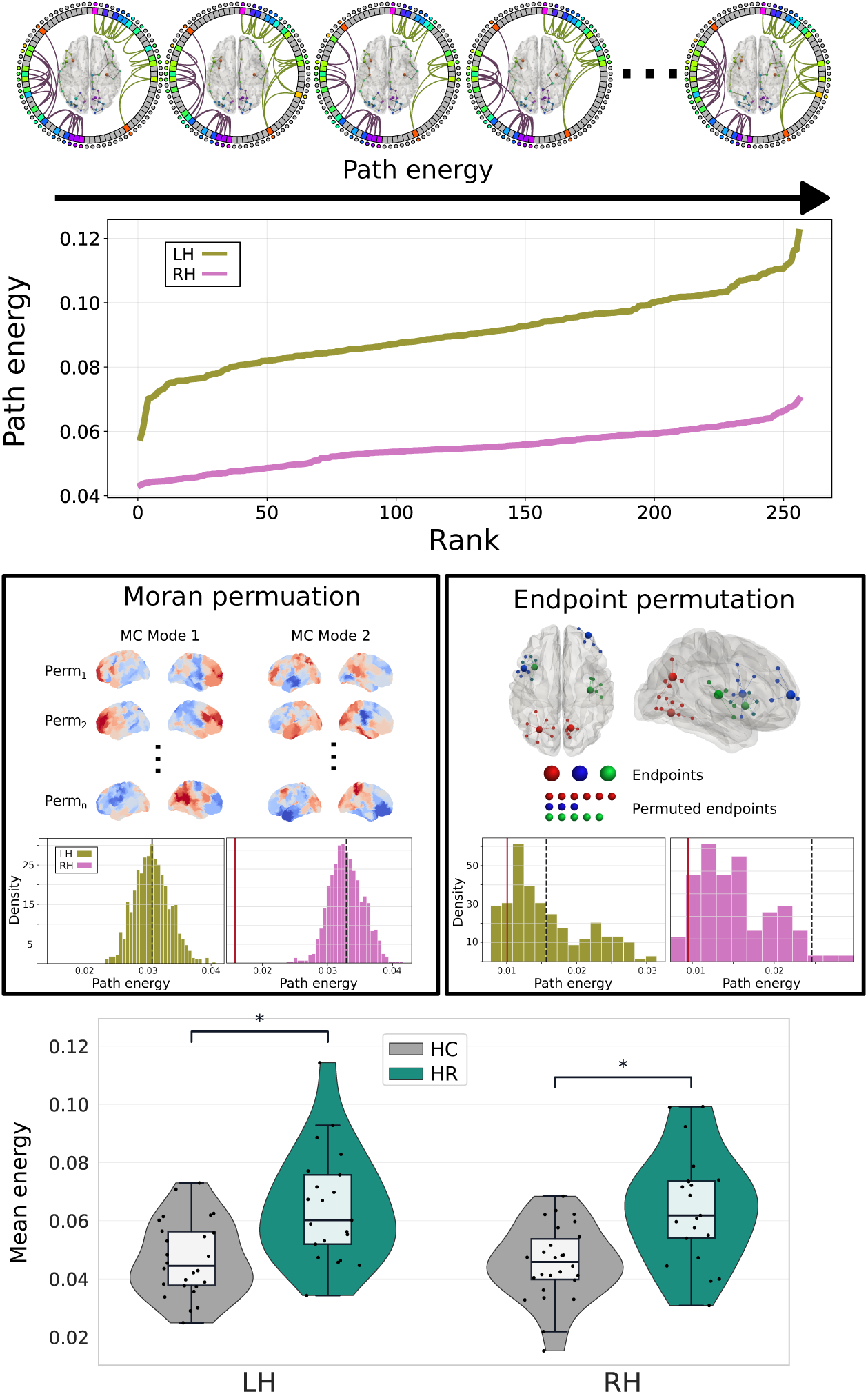
Path-ensemble construction and group differences in metabolic path energy. **Top**, Grey-matter-adjacent paths were constructed between subject-specific endpoints in the two-dimensional MC-mode space and ranked by segment-normalized path energy. Path energy increases with the magnitude of successive MC-mode displacements and their misalignment with the endpoint-defined direction. **Middle**, Moran-randomized MC modes and adjacent-endpoint perturbations used to generate the two path-energy null distributions. **Bottom**, Sex- and age-adjusted subject-level path energy in the left- and right-hemisphere cortical trajectories. Violin plots show participant distributions and points denote individual participants; brackets report the one-sided group-label permutation tests. Spatial-null probabilities are reported in the text.

## Discussion

In this study, we constructed matched metabolic, structural, and functional connectomes in HR individuals and healthy controls. Three main findings emerged. First, metabolic connectivity (MC) identified a distributed subnetwork with higher connectivity in HR, whereas no corresponding group-level differences were detected in structural connectivity (SC) or functional connectivity (FC). Contrary to our second hypothesis, the affected connections did not consistently converge on structural hubs: their nodes had modestly higher structural strength, but hub-node enrichment was absent in both hemispheres, edge-class enrichment was limited to feeder edges with parcellation-dependent lateralization, and hub–hub enrichment was not detected. Instead, the altered connections spanned the whole network, from short- to medium-range parcel pairs, and linked mostly non-adjacent regions that are weakly similar in controls but markedly more similar in HR. Second, parcel-wise concentrations of the individual metabolites contributing to MC did not differ significantly between groups after correction for multiple comparisons. The group effect therefore emerged primarily in the relationships between regional metabolic profiles rather than as focal changes in their absolute levels. Third, low-dimensional embedding revealed a reproducible metabolic organization extending from posterior unimodal to transmodal association cortex and, along a second cortical axis, from association to limbic territories. In controls, both axes were defined by antagonisms between metabolites rather than by their shared level: Ins and tNAA against Cho along the first, and Cho and tNAA against Glx along the second. HR participants showed reduced coherence of this organization, expressed as higher path energy along grey-matter-continuous trajectories through the MC-mode landscape.

### Metabolic connectivity alterations do not consistently converge on structural hubs

We hypothesized that the MC alteration would converge on structural hubs because hubs are topologically central and biologically costly (Bullmore and Sporns, 2012; van den Heuvel and Sporns, 2013), and because grey-matter lesions across brain disorders, including schizophrenia, are concentrated in hubs of the normative connectome (Crossley et al., 2014). The present data did not support this prediction. NBS nodes had modestly higher structural strength, but hub-node enrichment was absent in both hemispheres, feeder-edge enrichment appeared in only one hemisphere, which differed between parcellations, and hub–hub enrichment was not detected. The strength association is therefore better interpreted as a graded tendency of the altered subnetwork to include better-connected regions than as evidence of hub-specific vulnerability.

Several factors may explain this negative result. First, hub involvement may depend on illness stage. Rich-club and hub-related abnormalities have been reported in established schizophrenia (van den Heuvel et al., 2013; Klauser et al., 2017b), whereas evidence in clinical high-risk cohorts is mixed: structural rich-club organization was altered in one HR sample (Schmidt et al., 2017), while another study found no connectome abnormalities in the clinical high-risk for psychosis group overall, and fewer rich-club connections only among individuals who subsequently transitioned to psychosis (Hua et al., 2023). Because most individuals with HR do not transition, a hub effect restricted to those who will later convert could be diluted at the group level, although the present cross-sectional analysis cannot test this possibility. The absence of a significant SC component in the present sample is similarly not necessarily inconsistent with previous work, but emphasizes heterogeneity across high-risk populations.

Second, the evidence motivating our hypothesis derives from different measures. Hub vulnerability has been established primarily for grey-matter lesions and structural connections, whereas MC quantifies the similarity of multimetabolite profiles between regions. An MC edge need not correspond to a direct anatomical connection, so hubs of the structural connectome need not be the principal locus of altered metabolic similarity. Moreover, the metabolites contributing to MC are not direct readouts of the energetic demands that are central to accounts of hub vulnerability.

Third, the present sample was modest, and the strictest hub definitions contained few nodes, limiting sensitivity to hub-specific effects. In addition, NBS inference applies to the component as a whole rather than to its individual nodes or edges. The present results therefore do not exclude hub involvement in psychosis risk; rather, they indicate that the MC alteration was not reliably hub-centered in this sample and motivate examination of its distributed, large-scale organization.

The dissociation between metabolic and structural or functional group effects may be important for interpreting early stages of psychosis risk. The present cross-sectional data cannot establish whether metabolic alterations precede structural or functional dysconnectivity. Nevertheless, they show that abnormalities in the metabolic relationships between brain regions can be detectable in the absence of significant group differences in the corresponding structural and functional connectomes.

### Higher metabolic similarity between normally distinct regions

In controls, metabolic similarity decreased with the distance between regions, most clearly within hemispheres. This mirrors a general property of brain organization: inter-areal connection strength decays with distance (Ercsey-Ravasz et al., 2013), the brain’s spatial embedding contributes substantially to connectome topology (Roberts et al., 2016), and gene co-expression likewise decreases with distance (Arnatkeviciute et al., 2019). Part of the short-range similarity in MRSI is also expected to be technical, because the point-spread function and partial-volume effects blur metabolite signals across neighbouring parcels. For this reason, adjacent parcel pairs were distinguished in the edge-length analysis.

Against this background, the altered MC connections spanned the whole network, from short- to medium-range parcel pairs. They linked mostly non-adjacent regions, typically separated by a few intervening parcels along grey matter, whose metabolic profiles are weakly similar in controls, consistent with their separation, yet were markedly more similar in HR. Several considerations predict that such regions should remain metabolically distinct. Distant regions typically support different functions, from sensorimotor processing to transmodal association, and differ in neurotransmitter receptor and transporter distributions (Hansen et al., 2022) and in energy metabolism (Vaishnavi et al., 2010). They also occupy different positions along the S–A axis, along which cortical regions progressively diverge in microstructure, maturation timing, and connectivity during development (Sydnor et al., 2021; Paquola et al., 2019a). Association networks, in particular, are thought to have become untethered from the molecular gradients and early activity cascades that organize sensory hierarchies (Buckner and Krienen, 2013). Higher MC between such regions in HR may therefore reflect reduced metabolic differentiation of regions that normally diverge, in line with the reduced coherence of the metabolic gradient discussed below. Because MC is a statistical similarity measure and the present data are cross-sectional, whether this reflects altered developmental differentiation or ongoing physiological dysregulation remains to be determined.

### From regional metabolite abnormalities to metabolic network organization

Previous proton magnetic resonance spectroscopy studies in individuals at high risk for psychosis have primarily tested metabolite concentrations within predefined regions. Meta-analytic evidence has implicated several regional abnormalities, including altered glutamatergic, creatine, myo-inositol, and N-acetylaspartate measures in prefrontal, hippocampal, and thalamic regions, although findings vary considerably across studies, metabolites, and anatomical locations (Wang et al., 2020; Romeo et al., 2020; Whitehurst et al., 2020). Against this heterogeneous literature, we found no parcel in which an individual metabolite differed significantly between HR and controls after correction across the whole brain. Using voxel-based whole-brain MRSI rather than single-voxel MRS, a recent study from our group, in youth at risk for psychosis, reported higher frontal grey-matter tNAA in the pooled HR group and Ins elevations in the schizotypal subgroup, in the absence of structural or diffusion differences (Céléreau et al., 2026). Those effects were tested on absolute levels of individual metabolites, and their partial divergence from the present parcel-wise results may reflect differences in spatial resolution and inference (voxel-wise cluster statistics versus parcel averages). The present gradient analysis points to a complementary dimension: in controls, the MC modes were organized by antagonisms between metabolites rather than by their shared levels (Supplementary Tables S4 and S5). An alteration in the relative balance of metabolites across regions could therefore disrupt this organization without producing a detectable shift in any single metabolite.

The absence of focal level differences does not imply an absence of metabolic alteration. MC measures a different property: the similarity of multimetabolite profiles between regions. Two regions may therefore change their relationship to one another even when neither shows a sufficiently large univariate concentration shift to distinguish groups independently. In this sense, the present findings suggest that metabolic alterations associated with psychosis risk may be expressed partly as a reorganization of inter-regional metabolic relationships rather than solely as local increases or decreases in individual metabolites.

This distinction also provides a possible explanation for why MC revealed a significant group effect when SC and FC did not. MC samples a biochemical dimension of brain organization that is not reducible to either anatomical wiring or temporal BOLD coupling. The present findings therefore support a network-level extension of conventional MRS analysis, in which the organization of metabolic profiles across the brain becomes itself a phenotype of interest. This interpretation should nevertheless remain statistical: MC quantifies similarity between regional metabolic profiles and does not imply direct metabolite exchange or causal metabolic coupling between connected regions.

### Metabolic connectivity recapitulates cortical organizational gradients

A second major observation was that MC was not distributed arbitrarily across the cortex. Diffusion embedding revealed a dominant axis extending from posterior unimodal cortex, including visual regions, toward frontal transmodal association cortex. A second cortical progression extended from transmodal toward limbic and insular territories. The dominant axis closely resembles the sensorimotor-to-association (S–A) hierarchy previously described using functional connectivity, cortical microstructure, myelination, gene expression, and developmental imaging (Margulies et al., 2016; Sydnor et al., 2021; Paquola et al., 2019b; Whitaker et al., 2016; Paquola et al., 2019a).

The convergence is notable because the present gradient is derived from an entirely different signal domain: similarity between regional MRSI-derived metabolic profiles. It therefore suggests that the same macroscale hierarchy that structures anatomical and functional properties of the cortex is also expressed in its biochemical organization. The additional association-to-limbic progression indicates that metabolic organization is not reducible to a single rostro-caudal dimension, but occupies a low-dimensional manifold linking several major cortical territories.

The developmental significance of the dominant axis is particularly relevant to psychosis risk (Holmes et al., 2023). Cortical maturation proceeds hierarchically along the S–A axis, with primary sensory and motor systems maturing earlier and transmodal association territories showing more prolonged developmental trajectories (Sydnor et al., 2021; Whitaker et al., 2016; Gogtay et al., 2004). Intracortical myeloarchitectural changes during adolescence progressively differentiate higher-order from lower-order systems (Paquola et al., 2019a), while white-matter microstructure continues to mature into early adulthood (Lebel and Deoni, 2018). Intrinsic cortical activity follows the same developmental ordering: its maturational trajectories unfold along the S–A axis through late adolescence and are coupled to intracortical myelin maturation (Sydnor et al., 2023). Consistent with this organization, functional-connectivity refinement from childhood through adolescence varies systematically along the S–A axis and progressively reinforces the cortical hierarchy (Luo et al., 2024). Recent multispecies work further suggests that this hierarchy is actively established by opposing molecular developmental programs that differentiate primary sensorimotor from higher-order association cortex (Tsyporin et al., 2026). The spatial organization observed here may therefore reflect, at least in part, a metabolic expression of a cortical hierarchy whose origins are deeply developmental.

This interpretation does not imply that the adult MC gradient provides a direct record of developmental history. Rather, it motivates the hypothesis that the integrity of a spatial organization established through coordinated development may carry information about how coherently that organization has developed or is currently maintained.

### Reduced metabolic-gradient coherence in HR

This hypothesis motivated our path-based analysis. Visual inspection of the group-average MC-mode maps suggested that successive isolevels followed a more orderly spatial progression in controls than in HR. Because the cortical surface does not impose a unique linear ordering between the endpoint territories, we quantified this property across the set of anatomically admissible grey-matter-continuous paths rather than selecting a single parcel sequence.

Path energy measures how smoothly and directionally consistently such paths progress through MC-mode space. HR participants showed higher segment-normalized path energy than controls bilaterally, indicating a less coherent progression through the cortical metabolic landscape. The group differences remained significant after adjustment for age and sex and correction across hemispheres. The left-hemisphere effect was additionally supported by both spatial-autocorrelation and endpoint-randomization null models, whereas the right-hemisphere effect was robust to the spatial null but showed weaker robustness to perturbation of the endpoint locations. These controls are important because they indicate that the observed effect cannot be attributed solely to the strong spatial autocorrelation inherent to cortical gradients, while also showing some dependence on the anatomical definition of the right-hemisphere endpoints.

The path result provides a complementary level of description to the NBS finding. NBS identifies specific pairwise relationships whose MC strength differs between groups; path energy instead asks whether the larger spatial organization formed by those relationships remains coherent along the principal metabolic axes. Thus, the two analyses capture different manifestations of the same broader phenomenon: one localizes altered connections, whereas the other quantifies the integrity of the macroscale metabolic organization in which those connections are embedded.

This distinction may also help reconcile the present findings with previous gradient studies in psychosis. Gradient analyses have identified altered hierarchical organization across cortical, cerebellar, and subcortical functional connectomes in schizophrenia, including drug-naive first-episode illness, while related compression of the functional hierarchy has also been associated with schizotypy (Dong et al., 2020, 2023; Yang et al., 2023; Dong et al., 2025). Evidence at earlier illness stages is nevertheless inconsistent. In particular, no significant alterations of the primary or secondary functional gradients were detected in early psychosis, whereas established schizophrenia showed altered organization of the secondary visual-to-sensorimotor gradient; the primary S–A gradient remained comparatively preserved in both cohorts (Holmes et al., 2023). Our findings do not directly contradict these observations because metabolic and functional gradients quantify different physiological domains and because our path-energy measure assesses spatial coherence rather than regional gradient position or dispersion. Instead, they raise the possibility that biochemical organization along the cortical hierarchy may be altered at a stage at which corresponding macroscale structural or functional differences are not readily detected.

### Neurodevelopmental interpretation and limitations

The correspondence between the dominant MC axis and the S–A hierarchy provides a potential neurodevelopmental interpretation of reduced gradient coherence in HR. If the adult metabolic landscape is partly shaped by the same coordinated developmental processes that establish cortical hierarchy, then a less orderly progression along this axis could reflect altered establishment or maintenance of this organization. This interpretation is consistent with neurodevelopmental models of psychosis (Murray and Lewis, 1987; Insel, 2010), particularly given the prolonged maturation and plasticity of association cortex during adolescence and early adulthood (Sydnor et al., 2021; Whitaker et al., 2016).

The present study, however, cannot determine whether reduced metabolic-gradient coherence originates during development. The data are cross-sectional, and the same phenotype could result from earlier developmental perturbation, ongoing physiological dysregulation, or their interaction. Longitudinal studies following individuals across adolescence and through clinical outcomes will therefore be required to establish whether MC-gradient coherence changes over time or predicts transition to psychosis. Such designs can also separate trait-like from state-dependent neural features, and the present group-level contrast will need to be complemented by analyses relating inter-individual variation in MC-gradient coherence to symptoms, cognition, and functional outcome before it can inform patient stratification (Anticevic and Yang, 2026).

Several additional limitations should be considered. The path analysis depends on the definition of endpoint territories and on the chosen parcellation and grey-matter adjacency representation, although the use of alternative parcellations and spatial and endpoint null models mitigates some of these concerns. Our MC and path analyses were restricted to grey-matter parcels and therefore did not examine white-matter metabolic organization. White matter may nevertheless contain complementary information and future studies should determine whether related metabolic alterations extend into white matter and whether grey-white matter metabolic organization contributes to HR-related dysconnectivity. Sex and age were included as covariates in all group comparisons, but their specific influence on metabolic-gradient coherence was not examined. Sex differences in cortical organization are expressed along the S–A axis (Serio et al., 2024), and this axis varies with sex-hormone fluctuations within individuals (Serio et al., 2025). Because maturation along this axis continues into early adulthood (Sydnor et al., 2021), the age range of our cohort, larger samples will be needed to characterize how sex, age, and pubertal and hormonal status shape metabolic-gradient coherence and whether they moderate its alteration in HR. Finally, the modest sample size limits inference about clinical heterogeneity within HR, including whether the observed effects differ between individuals who subsequently remit, persist in a HR state, or transition to psychosis.

## Conclusion

Together, our findings identify a distributed alteration of metabolic network organization in HR that is not accompanied by detectable group-level structural or functional dysconnectivity. These metabolic differences do not consistently converge on structural hubs and are not explained by focal abnormalities in individual metabolite concentrations, highlighting the importance of the relationships between regional metabolic profiles. We further show that metabolic connectivity is organized along a low-dimensional cortical hierarchy resembling the sensorimotor-to-association axis and an association-to-limbic progression, and that this organization is less spatially coherent in HR. These results position whole-brain metabolic connectivity as a complementary dimension of connectomics and suggest that the coherence of macroscale metabolic organization may provide a systems-level phenotype of psychosis risk.

## Materials and methods

### Groups

#### Lausanne psychosis cohort

This study was conducted as part of the ongoing Lausanne Psychosis Cohort in Lausanne, Switzerland, and was approved by the local Cantonal Ethics Committee for Research on Human Beings (BASEC ID 2024-01777). Written informed consent was obtained from all adult participants. For minors, written consent was obtained from a parent or legal guardian, and written assent was obtained from the participant. HR participants were recruited through the Treatment and Early Intervention in Psychosis Program in Lausanne Baumann et al. (2013), which includes a dedicated early-detection pathway for individuals at risk for psychosis Conchon et al. (2023). HC participants were recruited in parallel and were selected to be comparable to the HR group in terms of age and sex distribution.

After restricting the cohort to healthy controls and HR participants, the pre-QC sample comprised 62 participants: 31 healthy controls (15 female / 16 male; age: 26.29 ± 7.10 years) and 31 HR participants (9 female / 22 male; age: 21.59 ± 4.15 years). Individuals in the HR group were assessed with the Structured Interview for Psychosis-Risk Syndromes (SIPS; ref). HR group comprised patients who met criteria for schizotypal personality disorder (SCZT) or attenuated psychosis syndrome (APS) according to Diagnostic and Statistical Manual of Mental Disorders, Fifth Edition, Text Revision Edition et al. (2013). APS and SCZT participants were pooled into a single High Risk for Psychosis (HR) group for all analyses.Healthy controls had no lifetime psychiatric diagnosis and no first-degree relative with a psychosis-related disorder.

Matrices entering modality-specific quality control were available for 31 healthy controls (15 female / 16 male; age: 26.29 ± 7.10 years) and 29 HR participants (9 female / 20 male; age: 21.90 ± 4.08 years). No additional diffusion-MRI dataset was excluded, so this was the final structural-connectivity sample. For metabolic connectivity, two HR participants were excluded because of poor raw MRSI quality and one control was excluded because of a sparse connectivity matrix, yielding 30 healthy controls (15 female / 15 male; age: 26.33 ± 7.21 years) and 27 HR participants (8 female / 19 male; age: 21.88 ± 4.24 years). For functional connectivity, seven datasets (four controls and three HR participants) were excluded because of severe motion artifacts in the source fMRI acquisitions, yielding 27 healthy controls (14 female / 13 male; age: 26.82 ± 7.42 years) and 26 HR participants (8 female / 18 male; age: 22.03 ± 4.28 years).

### MRI image acquisition

Magnetic resonance imaging (MRI) data were acquired at the CIBM Center for Biomedical Imaging, Lausanne University Hospital (CHUV), using a 3-Tesla scanner (MAGNETOM PrismaFit, Siemens Healthineers, Forchheim, Germany) equipped with a 32-channel head coil.

#### Anatomical imaging

High-resolution anatomical images were acquired using a T1-weighted magnetization-prepared rapid acquisition gradient echo (MPRAGE) sequence. Images were acquired with a voxel size of 1.0 × 1.0 × 1.2 mm^3^ and a matrix size of 240 × 256 × 160 voxels, corresponding to a field of view of 240 × 256 × 192 mm^3^. Sequence parameters were: repetition time (TR) = 2300 ms, echo time (TE) = 2.98 ms, and inversion time (TI) = 900 ms.

#### Diffusion-weighted imaging

Diffusion-weighted data were acquired using a diffusion spectrum imaging (DSI) sequence comprising 128 diffusion-weighted volumes with a maximum b-value of 8000 s/mm^2^, together with one non-diffusion-weighted b_0_ reference image. The DSI acquisition matrix was 96 × 96 × 34 voxels, with an isotropic voxel size of 2.2 mm. Sequence parameters were: TR = 6800 ms and TE = 144 ms.

#### Resting-state functional MRI

Resting-state functional MRI data were acquired with a repetition time of 1920 ms and an isotropic voxel size of 3.3 mm. Image quality was assessed using visual inspection and standard quality-control metrics to ensure data suitability for subsequent analyses.

#### Magnetic resonance spectroscopic imaging

Whole-brain ^1^H-MRSI data were acquired in a subset of participants from the Lausanne Psychosis Cohort. A first subcohort was scanned using a 3D ^1^H-FID-MRSI sequence and included 13 healthy controls (4 women / 9 men; age = 23.35 ± 3.51 years) and 21 HR participants (5 women / 16 men; age = 22.29 ± 4.20 years) before quality-control exclusions.

The sequence parameters were: echo time (TE) = 1.0 ms, repetition time (TR) = 353 ms, and flip angle = 40^◦^. The field of view and spatial resolution were matched to the Lausanne acquisition protocol, with a nominal voxel size of 5 × 5 × 5.3 mm^3^. Spectral bandwidth and free induction decay vector size were kept identical to the reference protocol.

A separate unsuppressed water acquisition was acquired for metabolite quantification and referencing. The water reference acquisition used TE = 1.07 ms, TR = 25 ms, and flip angle = 5^◦^. Field of view, spatial resolution, spectral bandwidth, and vector size were identical to those of the metabolite acquisition.

A secondary subcohort from the same recruitment stream was scanned using the ECCENTRIC 3D ^1^H-FID-MRSI sequence, a non-Cartesian spatial–spectral encoding approach based on 3D spherical stack-of-ECCENTRIC sampling and compressed-sensing reconstruction. The ECCENTRIC subcohort included 18 healthy controls (11 women / 7 men; age = 28.42 ± 8.29 years) and 10 HR participants (4 women / 6 men; age = 20.13 ± 3.81 years) before quality-control exclusions.

The ECCENTRIC acquisition used in the present cohort was performed with the following protocol-specific parameters. The metabolite acquisition had a field of view of 220 × 220 × 130 mm^3^ and a nominal voxel size of 5.0 × 5.0 × 5.2 mm^3^, corresponding to a scan resolution of 44 × 44 × 25. The sequence was acquired with TR = 457 ms, TE_1_ = 0.78 ms, TE_2_ = 65 ms, and flip angle = 45^◦^. The spectral bandwidth was 1320 Hz, with a vector size of 512 points and an acquisition duration of 389 ms. Spatial encoding was performed using an ECCENTRIC trajectory with an acceleration factor of 2.5 and an ECCENTRIC circle radius of 0.25 *k*_max_. The total acquisition time was 6 min 54 s.

A separate unsuppressed water reference acquisition was acquired with matched spatial coverage but lower spatial resolution. The water reference had a field of view of 220 × 220 × 130 mm^3^ and a nominal voxel size of 10.0 × 10.0 × 10.0 mm^3^, corresponding to a scan resolution of 22 × 22 × 13. The water reference sequence used TR = 460 ms, TE_1_ = 0.72 ms, TE_2_ = 65 ms, and flip angle = 45^◦^. The spectral bandwidth was 1320 Hz, with a vector size of 512 points and an acquisition duration of 389 ms. The water reference was acquired with water suppression turned off, an acceleration factor of 2.0, and an ECCENTRIC circle radius of 0.25 *k*_max_. The acquisition time for the water reference scan was 1 min 21 s. This acquisition was used for coil combination, field correction, and metabolite intensity normalization.

Both MRSI acquisitions were reconstructed using a compressed-sensing SENSE low-rank framework with total-generalized-variation regularization and simultaneous lipid suppression. Reconstructed spatio-spectral data were quantified voxel-wise using LCModel, with the water signal from the additional unsuppressed acquisition used as an internal reference. Because the acquisitions used ultra-short echo times, no explicit T2 relaxation correction was applied. Although the reduced TR introduces T1 weighting, only gray matter–gray matter metabolic connectivity was considered in the present study; therefore, explicit T1 correction would primarily introduce a global scaling factor and was not expected to alter the reported inter-regional similarity patterns. Post-quantification processing is described below.

The LCModel basis set included N-acetylaspartate (NAA), N-acetylaspartylglutamate (NAAG), creatine (Cr), phosphocreatine (PCr), glycerophosphocholine (GPC), phosphocholine (PCh), myoinositol (mI), scyllo-inositol (sI), glutamate (Glu), glutamine (Gln), lactate (Lac), gamma-aminobutyric acid (GABA), glutathione (GSH), taurine (Tau), aspartate (Asp), and alanine (Ala). Because only a subset of metabolites could be reliably resolved at the available field strength and spectral resolution, and because several compounds have overlapping resonances, metabolite estimates were combined into five robust metabolite volumes: total N-acetylaspartate (*t*NAA = NAA + NAAG), total creatine (*t*Cr = Cr + PCr), choline-containing compounds (Cho = GPC + PCh), myo-inositol (Ins), and glutamate plus glutamine (Glx). LCModel also provided voxel-wise spectral quality metrics, including signal-to-noise ratio (SNR) and Cramér–Rao lower bounds (CRLBs) for metabolite estimates.

### Preprocessing and connectome construction

All imaging-derived connectomes were reconstructed subject-wise using three matched volumetric parcellation schemes. These parcellations differed in their cortical subdivision but shared the same subcortical, cerebellar, and brainstem definitions, thereby allowing structural, functional, and metabolic connectivity to be compared within homologous nodal spaces.

#### Anatomical preprocessing and parcellation

For each participant, the T1-weighted anatomical image was processed with FreeSurfer version 7.2.0 (http://surfer.nmr.mgh.harvard.edu). The anatomical image was used to segment the brain and identify major supra-regions, including cortex, basal ganglia, thalamus, amygdala, hippocampus, hypothalamus, cerebellum, brainstem, and white matter.

Subject-specific volumetric parcellations were generated using the Chimera parcellation framework^77^. Three parcellation schemes were constructed for each participant and are hereafter referred to as LFMIHIFS-3, MFMIHIFS-197, and SFMIHIFS-200. These schemes differed in their principal cortical gray-matter parcellation:

- **L**: Lausanne cortical parcellation, scale 3^78^;
- **M**: MIST parcellation, 197 regions, covering cortical, subcortical, and cerebellar gray matter^79^;
- **S**: Schaefer cortical parcellation, 200 regions.

For the Lausanne- and Schaefer-based schemes, the cortical parcellation was combined with a common set of subject-specific subcortical and infratentorial parcellations intergated by Chimera:

- basal ganglia parcellation^80^;
- thalamus parcellation^81^;
- amygdala parcellation^82^;
- hippocampus parcellation^83^;
- hypothalamus parcellation^84^;
- brainstem parcellation^83^;
- cerebellar parcellation^85^.

The resulting subject-specific parcellation schemes comprised X parcels for LFMIHIFS-3, X parcels for MFMIHIFS-197, and X parcels for SFMIHIFS-200. For each participant, these three parcellation schemes were subsequently used as common nodal definitions for structural, functional, and metabolic connectome construction.

#### Structural connectivity

Diffusion spectrum imaging data were visually inspected for signal loss and motion-related artifacts that could compromise tractography quality^85^. DSI data were reconstructed following the approach described by Wedeen et al.^86^. Diffusion preprocessing, anatomically constrained tractography, and structural connectivity reconstruction were performed using MRtrix3^87^.

Peaks of the local fiber orientation distribution functions were estimated, with a maximum of three peaks retained per voxel, and used as input for deterministic tractography^88^. Tractography was performed with a maximum streamline length of 300 mm, 32 seeds per white-matter voxel per diffusion direction, and termination criteria based on either an angle greater than 60^◦^ between successive propagation steps or arrival at the white–gray matter boundary. For each participant, a tractogram of 10^6^ streamlines was retained to ensure reproducibility^89^.

For each of the three parcellation schemes, a weighted undirected structural connectivity matrix was reconstructed by assigning the whole-brain tractogram to the corresponding subject-specific regions. Streamline weights were refined using Spherical-deconvolution Informed Filtering of Tractograms (SIFT)^90^, which adjusts streamline contributions according to their consistency with the underlying diffusion signal. The nodes of each structural connectome corresponded to the regions of the LFMIHIFS-3, MFMIHIFS-197, or SFMIHIFS-200 parcellation.

The structural connection strength *S_ij_* between regions *i* and *j* was defined from the number of streamlines *ns_ij_* connecting the two regions, weighted by the inverse of their mean streamline length and corrected for regional volume. Length and volume corrections were applied to reduce tractography-related biases toward longer fibers and larger regions of interest, respectively^45^. To improve consistency across participants and reduce the influence of potential false-positive connections, edges present in fewer than 50% of participants were discarded^91,92^. This procedure yielded one structural connectivity matrix per participant and parcellation scheme.

#### Functional connectivity

Resting-state functional MRI data were preprocessed using fMRIPrep. Preprocessing included motion correction, slice-timing correction, co-registration to the anatomical image, spatial normalization, and nuisance signal regression. Anatomical information derived from CAT12 and FreeSurfer was used to support gray-matter masking, skull stripping, cortical surface reconstruction, and subcortical parcellation.

For each participant, regional resting-state fMRI time series were extracted separately using the LFMIHIFS-3, MFMIHIFS-197, and SFMIHIFS-200 parcellation schemes. Regional time-series quality was assessed at the parcel level, and parcels with insufficient spatial coverage or low BOLD signal reliability across participants were excluded prior to matrix construction. This exclusion mainly affected cerebellar regions, which were removed from the FC analyses, as well as parcels in areas prone to low signal-to-noise ratio or incomplete coverage, including the hypothalamus, amygdala, brainstem, and inferior temporal gyri. The retained subcortical node set therefore consisted primarily of thalamic nuclei, caudate, putamen, pallidum, and accumbens parcels.

The resulting regional time series were detrended and demeaned, and then band-pass filtered in the 0.04–0.07 Hz frequency range. Individual functional connectivity matrices were defined as the pairwise Pearson correlation coefficients between regional fMRI time series. This procedure yielded one functional connectivity matrix per participant and parcellation scheme.

### Metabolic connectivity

Post-quantification MRSI preprocessing was performed with MRSIprep Lucchetti et al. (2026), following the MC construction framework described previously Lucchetti et al. (2025). Briefly, MRSIprep registered quantified metabolite maps to each participant’s T1-weighted anatomy, generated tissue probability maps, applied voxel-wise partial-volume correction with a 5-mm point-spread-function model, projected the subject-specific Chimera parcellations into native MRSI space using nearest-neighbor interpolation, and generated standardized spatial-coverage and spectral-quality control measures. The LFMIHIFS-3, MFMIHIFS-197, and SFMIHIFS-200 parcellations were then applied to the five metabolite volumes. Parcels with inadequate gray-matter MRSI coverage or unreliable spectral quality were excluded before matrix construction; this principally affected susceptibility-prone orbitofrontal and inferior temporal regions, small amygdalar and hypothalamic parcels, and the cerebellum. The retained subcortical set consisted primarily of thalamic, caudate, putamen, pallidum, and accumbens parcels.

We use metabolic connectivity to denote an edge-wise relationship between regional representations over the metabolic signal domain. To obtain a robust and biochemically interpretable representation, MC was constructed from post-quantification metabolite maps rather than directly from raw spectra.

For every retained parcel, a regional metabolic profile was estimated by computing the median signal of each metabolite within that parcel. This yielded a five-dimensional metabolic profile per region:

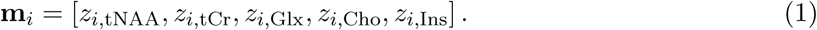

where metabolite values were z-score normalized across parcels within each participant and metabolite. This normalization removed global between-metabolite scaling differences and expressed each region according to its relative metabolic profile within the individual brain.

To account for uncertainty in metabolite quantification, regional metabolic profiles were augmented using a Monte Carlo uncertainty-propagation procedure. For each metabolite map, voxel-wise metabolite estimates were perturbed according to their LCModel-derived Cramér-Rao lower-bound variance, and this procedure was repeated (*K*_pert_ = 50) times. The perturbed profiles were used together with the original profile to stabilize the estimation of inter-regional metabolic connectivity.

For each participant and each parcellation scheme, the metabolic connectivity matrix (MC) was defined as the pairwise Spearman correlation between the metabolic profiles of all retained brain regions. Thus, each matrix element (*M_ij_*) quantified the similarity between the five-metabolite profiles of regions (i) and (j):

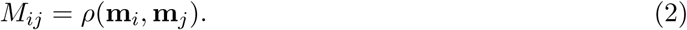

This procedure yielded one weighted, undirected MC matrix per participant for each of the three parcellation schemes: LFMIHIFS-3, MFMIHIFS-197, and SFMIHIFS-200.

#### Cross-sequence harmonization

Because MC matrices were derived from two MRSI acquisition protocols, namely the original 3D ^1^H-FID-MRSI sequence and the ECCENTRIC 3D ^1^H-FID-MRSI sequence, we harmonized individual metabolic connectivity matrices to reduce sequence-related effects prior to group-level analyses. Harmonization was performed separately for each parcellation scheme using neuroCombat, with MRSI sequence specified as the batch variable.

For each participant and parcellation scheme, the upper triangular elements of the MC were vectorized and entered into the harmonization model. Sex, age, and diagnostic group were included as biological covariates to be preserved during harmonization. Thus, neuroCombat estimated and removed additive and multiplicative effects associated with MRSI sequence while retaining variance attributable to age, sex, and diagnosis. The harmonized edge vectors were then reconstructed into symmetric MC matrices for subsequent analyses.

Formally, each edge-wise metabolic connectivity value was modeled as

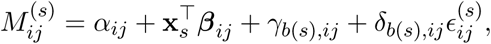

where 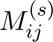 denotes the metabolic connectivity between regions *i* and *j* for subject *s*, **x***_s_* contains the preserved biological covariates, namely age, sex, and diagnostic group, and *b*(*s*) indexes the MRSI sequence used for subject *s*. Here, *γ_b_*_(_*_s_*_)_*_,ij_* captures sequence-specific additive shifts in the mean MC edge value, *δ_b_*_(_*_s_*_)_*_,ij_* captures sequence-specific multiplicative effects on edge-wise variance, and 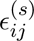 denotes residual subject-level variability not explained by the preserved biological covariates. Results of the harmonization are recapitulated in Supplementary Material (see REF).

### 0.1 Modality-specific matrix alignment and node inclusion

For each modality, connectivity matrices were constructed and analyzed separately using a modality-specific retained node set. Within each cohort and parcellation scheme, parcels were excluded from a given modality if valid regional profiles were absent or unreliable in more than 70% of participants. This ensured that, within each modality, all participant-level matrices were defined on a consistent set of nodes and could be compared across individuals without introducing edge-wise missingness. In particular, cerebellar parcels were entirely excluded from MC-based analyses due to insufficient MRSI coverage across participants. The resulting SC, FC, and MC matrices were therefore internally aligned within modality, but not forced onto a common cross-modal node set.

### Network-based analysis

Group differences in edge-wise connectivity were assessed using the network-based statistic (NBS), as introduced by Zalesky, Fornito, and Bullmore.^XX^ NBS is a non-parametric statistical framework designed to identify connected subnetworks showing significant group effects while controlling the family-wise error rate across the connectome. The method first performs mass-univariate testing at each edge, applies a primary test-statistic threshold, identifies connected components among suprathreshold edges, and then evaluates the significance of each component against a permutation-derived null distribution of maximal component size.

For each modality and parcellation scheme, individual connectivity or similarity matrices were entered into a MATLAB-compatible Python implementation of NBS. Edge-wise general linear models were fitted with diagnostic group as a categorical regressor of interest, sex as a categorical nuisance covariate, and age as a continuous nuisance covariate. The primary MC model additionally included MRSI acquisition sequence as a categorical nuisance covariate; a robustness model omitted this term. A primary edge-wise threshold was applied to the resulting test-statistic matrix, and connected components of suprathreshold edges were identified. Statistical significance was assessed using 5,000 Freedman–Lane permutations of reduced-model residuals, thereby retaining the nuisance-covariate structure while generating a null distribution of the largest connected component. Component-level *p*-values were computed as the proportion of permutations in which the maximal component size was greater than or equal to the observed component size. Positive and negative contrasts were tested separately; each directional component p-value was multiplied by two and capped at one to correct for the two tested directions. Components were considered significant at a two-direction-corrected network-level family-wise error rate of *p <* 0.05. All modality and MC robustness analyses used primary thresholds *t* = 3.00, 3.25, 3.50, 3.75, and 4.00. No additional correction across primary thresholds, parcellations, adjustment models, or modalities was applied.

### Grey matter adjacency

Parcels were considered adjacent if they shared a direct face-to-face boundary in the 3D label image, corresponding to 6-connectivity along the anterior–posterior, left–right, or inferior–superior voxel axes. For each non-zero parcel label, the label image was compared with shifted versions of itself along the six cardinal directions. Whenever two different non-zero parcel labels were observed in directly neighboring voxels, an undirected adjacency was assigned between the corresponding parcels. This yielded a binary adjacency matrix **A**^GM^, where

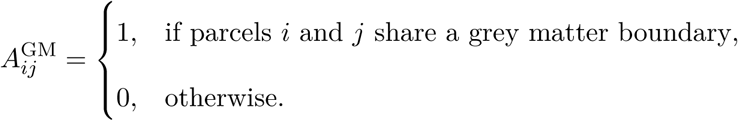

The matrix was symmetrized by construction and self-connections were excluded. Only labels included in the final parcellation were retained. When parcel names were available, direct cortex–cerebellum links were removed to avoid spurious adjacency induced by close spatial contact across non-contiguous anatomical compartments.

### Structural context of the metabolic NBS component

The significant sex- and age-adjusted MC NBS component at the primary threshold (*t* = 3.25) was mapped to the matching parcels of the structural connectome. An NBS node was defined as any parcel incident to at least one edge in the component, and its NBS load was the number of incident component edges. Structural connectivity matrices from healthy controls (Diag = 0) with matched MC data (*n* = 30) were averaged edge-wise (arithmetic mean) and symmetrized. Weighted structural node strength was then calculated for parcel *i* as

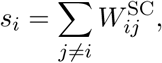

where *W* ^SC^ is the control-average structural connectivity matrix. The distributions of *s_i_* in NBS and non-NBS nodes were compared using a two-sided Mann–Whitney *U* test, with rank-biserial correlation reported as the effect size.

Hub and edge-class enrichment analyses were performed separately for each hemisphere. Structural hubs were defined independently at the top 2%, 5%, 10%, and 20% of the weighted node-strength distribution within each hemisphere; the top 10% definition was used for the principal structural map. At each threshold, the observed number of NBS nodes classified as hubs was compared with 5,000 node sets sampled uniformly without replacement from the same hemisphere, with every random set containing the same number of nodes as the hemisphere-specific set of NBS nodes. Enrichment was expressed as the observed overlap divided by the mean null overlap. The 2.5th and 97.5th percentiles of the null distribution defined its 95% interval, and upper-tail empirical probabilities were calculated with the finite-sample correction (*b* + 1)*/*(5,000 + 1), where *b* is the number of null overlaps at least as large as the observed overlap.

For the edge-level analysis, intrahemispheric NBS edges were classified as hub–hub, feeder, or local when both, one, or neither endpoint was a strength-defined hub, respectively. This classification was repeated at the 2%, 5%, 10%, and 20% hub thresholds. At each threshold, 5,000 sets containing the same number of edges as the hemisphere-specific NBS edge set were sampled without replacement from all possible undirected parcel pairs within that hemisphere. Interhemispheric NBS edges were not included in the edge-class analysis. For each edge class, enrichment was the observed count divided by its mean random-set count; 95% null intervals and upper-tail empirical probabilities were obtained from the corresponding random-set distribution.

Edge length was defined as the Euclidean distance between parcel centroids in world coordinates. Each NBS edge was additionally classified as adjacent or non-adjacent using the grey-matter adjacency matrix described above. For intrahemispheric NBS edges, the number of adjacency steps separating the two parcels was computed as the unweighted shortest-path length in the within-hemisphere grey-matter adjacency graph. The distance dependence of MC in controls was summarized descriptively as the Spearman correlation between control-average MC and edge length across all parcel pairs within each hemisphere.

### Parcel-wise metabolite analysis

To test whether the network-level MC findings could be attributed to focal differences in metabolite concentration, raw parcel values were analyzed separately for tNAA, tCr, Glx, Cho, and Ins. Before statistical testing, parcel-wise values were harmonized separately for each metabolite using neuroCombat. MRSI acquisition sequence was specified as the batch variable, while diagnosis, sex, and age were included as biological covariates to be preserved. neuroCombat estimated and removed sequence-associated additive and multiplicative effects without removing variance attributable to these prespecified covariates. Because neuroCombat requires complete features, parcels containing a non-finite value in any included participant were omitted from this analysis, leaving 245 parcels for each metabolite. For every retained metabolite–parcel pair, an ordinary least-squares model included diagnosis as the regressor of interest and sex and age as nuisance covariates,

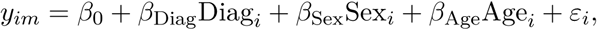

where *y_im_* is metabolite *m* in the parcel for participant *i*, controls were the reference group (Diag = 0), and HR participants were coded as Diag = 1. The diagnostic coefficient was evaluated with a two-sided *t* test using complete observations for the corresponding parcel–metabolite pair. Benjamini–Hochberg false-discovery-rate correction was applied across parcels separately within each metabolite, with *q <* 0.05 considered significant.

### Gradient analysis

#### Metabolic connectivity modes

MC modes were estimated with the BrainSpace diffusion-maps implementation Vos de Wael et al. (2020). Because MC matrices contained signed correlations, edge values were transformed from [−1, 1] to [0, 1] as *W* ^′^ = (*W* + 1)*/*2 and clipped to this interval. Affinity between parcel-wise MC profiles was computed using the normalized-angle kernel. The default sparsity parameter of 0.9 was used, removing the weakest 90% of affinities within each row before embedding. Diffusion maps were fitted with 10 components, anisotropic normalization *α* = 0.5, and diffusion time *t* = 0, corresponding to the multiscale diffusion-map embedding. No additional min–max normalization or coordinate scaling was applied (scale factor 1).

The same embedding procedure was applied independently to every subject-specific MC matrix and to the population-average MC matrix. The latter provided the common alignment reference. Each subject-level component was aligned separately to the corresponding reference component using BrainSpace generalized Procrustes alignment with 10 iterations and a convergence tolerance of 10^−5^. Alignment was performed without centering or scaling, thereby resolving component orientation and sign while preserving the estimated component amplitudes. Control-average coordinates used for the group-level visualizations were calculated by averaging the aligned control embeddings. The first two aligned MC modes retained for the triangular manifold and path analyses are denoted *g*_1_ and *g*_2_.

#### Principal gradient vectors

To summarize the geometry of the two-dimensional metabolic embedding, we fitted a reference triangle to the parcel point cloud in the space defined by the first two aligned MC modes (*g*_1_, *g*_2_). The convex hull of the point cloud was first computed, and the maximal-area triangle among candidate hull vertices was selected as a compact approximation of the embedding boundary. The three vertices of this triangle defined representative endpoints of the metabolic gradient manifold. They were ordered as the apex, left-base, and right-base vertices, corresponding to the three main directions of variation observed in the embedding.

The triangle is an empirical approximation rather than a constraint imposed by diffusion embedding. Its spectral motivation follows from an idealized network model in which MC favors strong local coupling along grey-matter-continuous geometry while retaining global edge diversity. In the continuum limit, the resulting spectral modes obey a Neumann eigenproblem; when the first two nonconstant modes share an intrinsic path coordinate, they satisfy the approximately parabolic relation 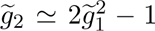. Anatomical heterogeneity and branching can broaden this curve into the triangular cloud observed empirically (Supplementary Mathematical Note S1).

For each edge of the fitted triangle, a principal gradient vector was defined as the unit vector connecting two endpoint vertices. Let **v***_a_* and **v***_b_* denote the two-dimensional coordinates of two triangle vertices. The corresponding principal gradient direction was defined as

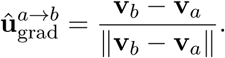

This yielded three principal gradient vectors, corresponding to the three edges of the triangular manifold. These vectors were used to quantify the alignment of network paths with each major axis of metabolic variation, including the unimodal-to-transmodal, transmodal-to-limbic, and transmodal-to-subcortical trajectories.

Principal trajectories could be defined either by a single triangle edge or by a piecewise sequence of edges passing through an intermediate vertex. For a trajectory composed of vertices (*a, b, c*), the corresponding direction was therefore treated as two directed branches, *a* → *b* and *b* → *c*, rather than as a single straight-line vector from *a* to *c*. This allowed curved trajectories on the embedding manifold to be represented as ordered combinations of elementary triangle-derived gradient directions.

#### Metabolite contrasts underlying the MC modes

To identify which metabolites drive each MC mode, we related *g*_1_ and *g*_2_ to the spatial distributions of the five metabolites (tCr, Glx, Cho, tNAA, and Ins). For each metabolite, a group-average map was constructed by averaging parcel-wise concentrations across healthy controls. Each group-average map was then *z*-scored across parcels, removing differences in concentration scale between metabolites. For every pair of metabolites *A* and *B*, two maps were derived:

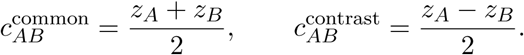

The common map captures variation in which the two metabolites co-vary in the same direction, whereas the contrast map captures variation in which one metabolite is elevated relative to the other. Both maps were correlated (Pearson *r*) with each control-average MC mode across parcels with valid data for both metabolites. A high |*r*| for the contrast map combined with a low |*r*| for the common map indicates that a mode reflects a relative difference between two metabolites, one rising as the other falls across the cortex, rather than their shared overall level, such as global MRSI signal intensity or grey-matter content. A positive contrast correlation indicates that the first-named metabolite of the pair is relatively elevated toward higher mode values. The analysis was restricted to controls to characterize the normative organization of the MC modes.

### Path-ensemble energy

#### Grey-matter-constrained path generation

To quantify MC-mode organization beyond regional effects, we modeled the two-dimensional embedding as a constrained path space on the brain network. For each subject–session and hemisphere, the triangle-derived R, G, and B endpoints and their saved traversal order were retained. A path P = (*ℓ*_0_*, ℓ*_1_*, …, ℓ_L_*) was defined as an ordered sequence of parcels in which every consecutive pair (*ℓ_j_*_−1_*, ℓ_j_*) was connected in the grey-matter adjacency graph **A**^GM^. Each parcel *ℓ_j_* was associated with a two-dimensional MC-mode coordinate,

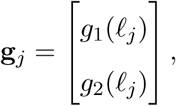

where *g*_1_ and *g*_2_ denote the first two aligned MC modes.

Paths were generated separately for each consecutive endpoint pair in the saved traversal order. Graph edges were weighted by Euclidean displacement in (*g*_1_*, g*_2_) space, and a weighted shortest path was first obtained for each pair. The maximum number of nodes permitted in a candidate path was the smaller of the shortest-path node count plus four, the number of available parcels, and 24; candidate paths therefore contained at most 24 nodes (23 edges). We enumerated node-simple paths, with no repeated parcel within a segment, retaining at most 200 paths for each endpoint pair. The third triangle endpoint was excluded while searching between the other two. Pairwise segments were then joined in the saved endpoint order, and combinations that repeated a parcel across branches were discarded, except at their shared endpoint. Up to 10,000 complete cortical paths were retained per subject–session and hemisphere; when fewer valid combinations existed, all available paths were retained. No directional pre-filter was applied during enumeration, so directional coherence entered the analysis through the energy function rather than through path acceptance. The optional line-proximity penalty was also disabled.

#### Path energy

The displacement between two successive nodes along a path was defined as

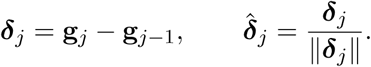

Path energy was defined from the geometry of the path in gradient space. Specifically, for each principal trajectory, we penalized path segments that deviated from the corresponding triangle-derived principal gradient vector 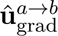:

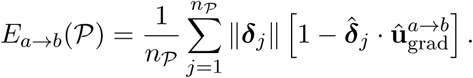

Here, the term 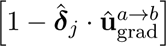 penalizes angular misalignment with the principal gradient direction, while the factor ∥***δ****_j_*∥, weights this penalty by the magnitude of the displacement in gradient space. Consequently, small deviations in gradient space contribute weakly, whereas large off-axis displacements contribute more strongly. The factor 1*/n*_P_ converts the cumulative cost into a mean segment-wise energy, reducing the trivial dependence of *E* on the number of graph steps in a path. This normalization allows paths with different numbers of anatomical segments to be compared in terms of their average directional efficiency in metabolic gradient space. Thus, paths composed of segments that advance consistently along the principal gradient incur lower energy, whereas tortuous, oppositely directed, or strongly off-axis trajectories incur higher energy.

For trajectories composed of multiple branches, path energy was computed as a globally normalized sum over the corresponding directed segments. Thus, for a piecewise trajectory passing through vertices (*a, b, c*), the total energy was defined as

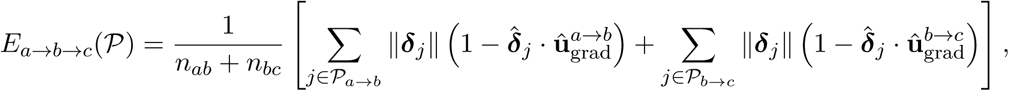

where *n_ab_* and *n_bc_* denote the number of anatomical segments in each branch. This preserves the interpretation of energy as a mean segment-wise directional cost, while allowing curved trajectories to be evaluated using branch-specific principal gradient vectors. More generally, for a piecewise trajectory composed of *M* directed branches, the same definition extends by summing the branch-specific costs and normalizing by the total number of anatomical segments across all branches. This formulation allows both straight and curved principal trajectories to be evaluated within the same path ensemble.

#### Subject-level path-ensemble energy

For each subject–session and hemisphere, every retained path 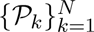 was assigned its segment-normalized energy *E_k_* = *E*(P*_k_*). To emphasize lower-energy paths while retaining all admissible configurations, normalized Boltzmann weights were defined as

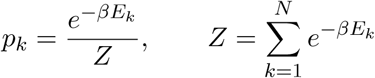

where *β* controls the relative weighting of lower-energy paths and was fixed to one in all analyses. The participant-level summary statistic was the weighted mean path energy,

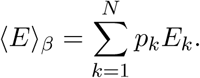

Lower ⟨*E*⟩*_β_* indicates that the retained paths advance through MC-mode space using smaller and more directionally aligned displacements; higher values indicate larger or more strongly misaligned steps. This weighted mean energy was used in the reported group analysis.

### Statistical analysis

#### Moran spatial null model

To test whether the observed path energy could be explained by spatial autocorrelation in the metabolic gradients, we generated spatially constrained surrogate gradient maps using Moran spectral randomization as implemented in BrainSpace Vos de Wael et al. (2020); Wagner and Dray (2015). Spatial weights were defined from inverse Euclidean distances between parcel centroids. The first two MC modes (*g*_1_ and *g*_2_) were randomized separately and paired to form 1000 surrogate two-dimensional embeddings for each participant. For every surrogate embedding, the grey-matter adjacency graph, selected endpoints, endpoint traversal order, and saved anatomical paths were held fixed. Individual path energies and the participant-level weighted mean energy were then recomputed using the same parameters as in the observed analysis, producing a participant-specific spatial null distribution of path energy. This procedure preserves the spatial autocorrelation of each MC mode while disrupting its observed anatomical arrangement.

#### Adjacent endpoint randomization

To assess whether path-energy estimates depended on the precise anatomical placement of the selected gradient endpoints, we generated a second null distribution by locally perturbing the endpoints while preserving grey-matter topology. For each R, G, and B endpoint, candidate replacements were restricted to immediate neighbors in the grey-matter adjacency graph within the corresponding hemisphere. The original endpoint nodes and candidates with invalid gradient co-ordinates were excluded. Let N*_R_*, N*_G_*, and N*_B_* denote the resulting candidate sets, with sizes *n_R_*, *n_G_*, and *n_B_*. Because the three endpoint identities remained fixed, the candidate configurations were the Cartesian product

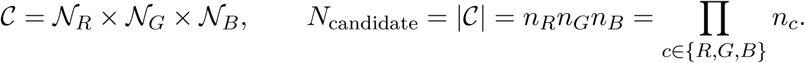

Thus, if each endpoint had *q* eligible neighbors, the procedure yielded *q*^3^ candidate triplets, rather than 3! permutations, because R, G, and B retained their labels and traversal order. Candidate triplets in which two endpoints mapped to the same parcel, or for which no valid complete grey-matter path could be generated, were discarded. Accordingly, the number entering the null distribution was

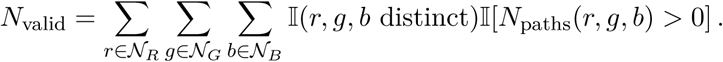

For each valid combination, paths were generated anew in the observed gradient embedding and the weighted mean path energy was recomputed using the same endpoint order, graph constraints, path-search limits, and energy parameters as in the primary analysis. The number of valid endpoint configurations was allowed to vary across participants because it depended on local graph degree and path availability. This procedure perturbs endpoint definition without randomizing the underlying metabolic gradient coordinates or introducing anatomically disconnected configurations.

#### Path-energy significance testing

Path-energy group comparisons were performed separately for left- and right-hemisphere cortical trajectories. Values were adjusted for sex and age using ordinary least squares, and healthy controls and HR participants were compared using a one-sided permutation test on the difference in group medians, with the alternative hypothesis of lower path energy in controls. Statistical significance of the observed group difference was estimated from 10,000 permutations of diagnostic labels.

The group comparison was additionally evaluated against the Moran and adjacent-endpoint path-energy distributions. For each null replicate, participant-level null energies were assembled across groups, adjusted for sex and age, and the same median-difference statistic was recomputed. Because the number of adjacent-endpoint configurations varied between participants, endpoint distributions were truncated to the minimum number available within each hemispheric analysis. Empirical lower-tail probabilities were calculated by comparing the observed group statistic with the corresponding null distribution. Moran and adjacent-endpoint probabilities were computed independently, and when both models were applied, the more conservative of the two probabilities was reported. The two null distributions were therefore not pooled or applied sequentially; significance required the observed group difference to remain supported under both spatial and endpoint uncertainty.

### Visualization

#### Triangular RGB Mapping of Gradient Embeddings

The same fitted triangle was also used for visualization. Each triangle vertex was assigned an RGB basis color, and each parcel was colored according to its barycentric coordinates relative to the fitted triangle. This produced a continuous three-way color mixture across the embedding, rather than a hard assignment to the nearest vertex. Barycentric weights were clipped to the valid interval, renormalized to sum to one, and the resulting RGB values were chroma-normalized to preserve hue while ensuring comparable saturation across parcels.

## Funding and Acknowledgments

This work was supported by the Swiss National Science Foundation (SNF) (grant number 215728). EC was supported by an MD-PhD fellowship from the Faculty of Biology and Medicine, University of Lausanne. PK was supported by a fellowship from the Adrian & Simone Frutiger Foundation. Recruitment for the Lausanne Psychosis Cohort was supported by SNF NCCR-Synapsy (grant number 185897) and the NeuroNA Foundation. The funders had no role in study design, data collection and analysis, decision to publish, or preparation of the manuscript.

## Competing interests

Antoine Klauser is employed by Siemens Healthcare as a research scientist. He contributes to on-site research and development initiatives. He does not hold fiduciary responsibilities for Siemens Healthcare. All the other authors declare no competing interests.

