## Supplementary Material for "Whole-Brain MRSI Metabolic Connectomics Reveals Network Disruptions and Altered Gradient Coherence in Psychosis Risk"

Metabolic connectivity disruptions in individuals at High Risk for Psychosis

#### NBS results across modalities and parcellations

Supplementary Table S1 reports the network-based statistic (NBS) results for metabolic (MC), functional (FC), and structural (SC) connectivity across the three Chimera parcellations (LFMIHIFIS-3, MFMIHIFIS-197, SFMIHIFIS-200) and five primary thresholds ($t_0$ = 3.00–4.00). All models used diagnostic group as regressor with sex and age as nuisance covariates; for MC, a sensitivity model additionally adjusted for MRSI sequence. Each run used 5,000 Freedman–Lane permutations and component extent, and is represented by its minimum-p component. P-values are corrected for the two tested directions but not across thresholds, parcellations, or modalities; "n/a" denotes runs in which no edge exceeded $|t_0|$. HC and HR values are the subject-wise mean connectivity over the component's edges. Samples after quality control were 30 HC/27 HR for MC, 27 HC/24 HR for FC, and 30–31 HC/28–29 HR for SC, depending on parcellation.

Table S1: **NBS results across modalities, parcellations, adjustment models, and primary thresholds.** One minimum-p component is shown per run. Component 0 with effect *none above* $|t_0|$ means that neither tested tail contained an observed edge beyond the primary threshold; because no component exists, its component-level p-value is not applicable. The observed test-statistic matrix of every row was independently reproduced under correct subject/session pairing and its declared nuisance model. HC and ARMS mean ± SD are computed over the edges of the selected component at each row's primary threshold $t_0$, so these values are not comparable across rows for the same modality and parcellation.

| Mod. | Chimera parcellation | Adjustment | $t_0$ | Comp. | Tail | Effect | $N$ HC/ARMS | Nodes/edges | HC mean ± SD | ARMS mean ± SD | $t$ mean ± SD | $p_{\text{saved}}$ |
|---|---|---|---|---|---|---|---|---|---|---|---|---|
| **Metabolic connectivity (MC)** | | | | | | | | | | | | |
| MC | LFMIHIFIS-3 | MRSI sequence + sex + age | 3.00 | 7 | left | ARMS > CTRL | 30/27 | 92/188 | 0.1129 ± 0.166 | 0.3801 ± 0.1559 | −3.442 ± 0.4291 | 0.0352 |
| MC | LFMIHIFIS-3 | MRSI sequence + sex + age | 3.25 | 5 | left | ARMS > CTRL | 30/27 | 66/109 | 0.1123 ± 0.1767 | 0.3975 ± 0.1694 | −3.675 ± 0.417 | 0.0272 |
| MC | LFMIHIFIS-3 | MRSI sequence + sex + age | 3.50 | 3 | left | ARMS > CTRL | 30/27 | 38/54 | 0.1348 ± 0.1832 | 0.4381 ± 0.1865 | −3.973 ± 0.4004 | 0.028 |
| MC | LFMIHIFIS-3 | MRSI sequence + sex + age | 3.75 | 2 | left | ARMS > CTRL | 30/27 | 33/37 | 0.1046 ± 0.1866 | 0.4216 ± 0.1946 | −4.139 ± 0.3798 | 0.012 |
| MC | LFMIHIFIS-3 | MRSI sequence + sex + age | 4.00 | 2 | left | ARMS > CTRL | 30/27 | 19/19 | 0.1132 ± 0.1712 | 0.4299 ± 0.191 | −4.42 ± 0.3348 | 0.0108 |
| MC | LFMIHIFIS-3 | Sex + age | 3.00 | 8 | left | ARMS > CTRL | 30/27 | 90/213 | 0.1324 ± 0.1653 | 0.397 ± 0.1509 | −3.443 ± 0.435 | 0.0284 |
| MC | LFMIHIFIS-3 | Sex + age | 3.25 | 3 | left | ARMS > CTRL | 30/27 | 68/120 | 0.1214 ± 0.1735 | 0.4033 ± 0.1683 | −3.694 ± 0.4197 | 0.026 |
| MC | LFMIHIFIS-3 | Sex + age | 3.50 | 3 | left | ARMS > CTRL | 30/27 | 47/67 | 0.1354 ± 0.1795 | 0.4328 ± 0.1757 | −3.95 ± 0.4027 | 0.0208 |
| MC | LFMIHIFIS-3 | Sex + age | 3.75 | 2 | left | ARMS > CTRL | 30/27 | 34/39 | 0.117 ± 0.1885 | 0.4311 ± 0.1925 | −4.185 ± 0.3699 | 0.0128 |
| MC | LFMIHIFIS-3 | Sex + age | 4.00 | 2 | left | ARMS > CTRL | 30/27 | 22/24 | 0.1082 ± 0.1748 | 0.4249 ± 0.199 | −4.393 ± 0.3274 | 0.0068 |
| MC | MFMIHIFIS-197 | MRSI sequence + sex + age | 3.00 | 6 | left | ARMS > CTRL | 30/27 | 60/96 | 0.1898 ± 0.1772 | 0.4536 ± 0.147 | −3.365 ± 0.3306 | 0.054 |
| MC | MFMIHIFIS-197 | MRSI sequence + sex + age | 3.25 | 9 | left | ARMS > CTRL | 30/27 | 37/47 | 0.203 ± 0.1788 | 0.4891 ± 0.1582 | −3.603 ± 0.3085 | 0.0544 |
| MC | MFMIHIFIS-197 | MRSI sequence + sex + age | 3.50 | 7 | left | ARMS > CTRL | 30/27 | 25/26 | 0.1699 ± 0.1849 | 0.4684 ± 0.1624 | −3.805 ± 0.2761 | 0.0404 |
| MC | MFMIHIFIS-197 | MRSI sequence + sex + age | 3.75 | 9 | left | ARMS > CTRL | 30/27 | 7/6 | 0.1803 ± 0.2307 | 0.4889 ± 0.2182 | −4.091 ± 0.2953 | 0.1196 |
| MC | MFMIHIFIS-197 | MRSI sequence + sex + age | 4.00 | 1 | right | CTRL > ARMS | 30/27 | 2/1 | 0.6532 ± 0.155 | 0.3125 ± 0.4437 | 4.01 | 0.7952 |
| MC | MFMIHIFIS-197 | Sex + age | 3.00 | 6 | left | ARMS > CTRL | 30/27 | 60/107 | 0.1968 ± 0.1811 | 0.4586 ± 0.1492 | −3.392 ± 0.341 | 0.042 |
| MC | MFMIHIFIS-197 | Sex + age | 3.25 | 9 | left | ARMS > CTRL | 30/27 | 41/54 | 0.2109 ± 0.1741 | 0.4938 ± 0.1506 | −3.626 ± 0.3174 | 0.0456 |
| MC | MFMIHIFIS-197 | Sex + age | 3.50 | 6 | left | ARMS > CTRL | 30/27 | 26/27 | 0.1841 ± 0.1818 | 0.4844 ± 0.1672 | −3.861 ± 0.2933 | 0.038 |
| MC | MFMIHIFIS-197 | Sex + age | 3.75 | 6 | left | ARMS > CTRL | 30/27 | 9/9 | 0.1815 ± 0.2259 | 0.4877 ± 0.2267 | −4.049 ± 0.3229 | 0.0676 |
| MC | MFMIHIFIS-197 | Sex + age | 4.00 | 1 | right | CTRL > ARMS | 30/27 | 2/1 | 0.6532 ± 0.155 | 0.3125 ± 0.4437 | 4.037 | 0.8036 |
| MC | SFMIHIFIS-200 | MRSI sequence + sex + age | 3.00 | 7 | left | ARMS > CTRL | 30/27 | 102/218 | 0.1309 ± 0.1739 | 0.3987 ± 0.1554 | −3.402 ± 0.3569 | 0.0312 |
| MC | SFMIHIFIS-200 | MRSI sequence + sex + age | 3.25 | 6 | left | ARMS > CTRL | 30/27 | 77/122 | 0.1108 ± 0.1765 | 0.39 ± 0.1571 | −3.629 ± 0.3268 | 0.028 |
| MC | SFMIHIFIS-200 | MRSI sequence + sex + age | 3.50 | 6 | left | ARMS > CTRL | 30/27 | 43/64 | 0.1603 ± 0.1949 | 0.4629 ± 0.1694 | −3.826 ± 0.317 | 0.0272 |
| MC | SFMIHIFIS-200 | MRSI sequence + sex + age | 3.75 | 5 | left | ARMS > CTRL | 30/27 | 27/31 | 0.1418 ± 0.1993 | 0.4608 ± 0.1721 | −4.064 ± 0.3013 | 0.0208 |
| MC | SFMIHIFIS-200 | MRSI sequence + sex + age | 4.00 | 4 | left | ARMS > CTRL | 30/27 | 12/11 | 0.1297 ± 0.2264 | 0.4748 ± 0.1869 | −4.256 ± 0.322 | 0.03 |
| MC | SFMIHIFIS-200 | Sex + age | 3.00 | 7 | left | ARMS > CTRL | 30/27 | 104/248 | 0.143 ± 0.1716 | 0.4088 ± 0.1552 | −3.411 ± 0.3717 | 0.0244 |
| MC | SFMIHIFIS-200 | Sex + age | 3.25 | 6 | left | ARMS > CTRL | 30/27 | 79/136 | 0.1333 ± 0.1765 | 0.4125 ± 0.1593 | −3.64 ± 0.3481 | 0.0248 |
| MC | SFMIHIFIS-200 | Sex + age | 3.50 | 4 | left | ARMS > CTRL | 30/27 | 46/69 | 0.1607 ± 0.1937 | 0.4647 ± 0.1665 | −3.883 ± 0.3221 | 0.0252 |
| MC | SFMIHIFIS-200 | Sex + age | 3.75 | 6 | left | ARMS > CTRL | 30/27 | 30/38 | 0.1463 ± 0.2044 | 0.4629 ± 0.1684 | −4.081 ± 0.3021 | 0.0168 |
| MC | SFMIHIFIS-200 | Sex + age | 4.00 | 4 | left | ARMS > CTRL | 30/27 | 11/10 | 0.138 ± 0.2242 | 0.4851 ± 0.1914 | −4.377 ± 0.2804 | 0.042 |

**Supplementary Table S1 (continued)**

| Mod. | Chimera parcellation | Adjustment | $t_0$ | Comp. | Tail | Effect | $N$ HC/ARMS | Nodes/edges | HC mean ± SD | ARMS mean ± SD | $t$ mean ± SD | $p_{\text{saved}}$ |
|---|---|---|---|---|---|---|---|---|---|---|---|---|
| **Functional connectivity (FC)** | | | | | | | | | | | | |
| FC | LFMIHIFIS-3 | Sex + age | 3.00 | 1 | right | CTRL > ARMS | 27/24 | 101/152 | 0.3281 ± 0.1004 | 0.1616 ± 0.09383 | 3.325 ± 0.2643 | 0.1408 |
| FC | LFMIHIFIS-3 | Sex + age | 3.25 | 1 | right | CTRL > ARMS | 27/24 | 65/81 | 0.3381 ± 0.101 | 0.1632 ± 0.09692 | 3.513 ± 0.223 | 0.114 |
| FC | LFMIHIFIS-3 | Sex + age | 3.50 | 2 | right | CTRL > ARMS | 27/24 | 26/26 | 0.3576 ± 0.1119 | 0.1736 ± 0.1167 | 3.725 ± 0.1842 | 0.1416 |
| FC | LFMIHIFIS-3 | Sex + age | 3.75 | 2 | right | CTRL > ARMS | 27/24 | 5/4 | 0.2693 ± 0.1384 | 0.08165 ± 0.1033 | 3.822 ± 0.02662 | 0.324 |
| FC | LFMIHIFIS-3 | Sex + age | 4.00 | 1 | right | CTRL > ARMS | 27/24 | 2/1 | 0.5241 ± 0.1793 | 0.2768 ± 0.2415 | 4.117 | 0.8148 |
| FC | MFMIHIFIS-197 | Sex + age | 3.00 | 1 | right | CTRL > ARMS | 27/24 | 38/39 | 0.3897 ± 0.09529 | 0.226 ± 0.1003 | 3.291 ± 0.271 | 0.4192 |
| FC | MFMIHIFIS-197 | Sex + age | 3.25 | 2 | right | CTRL > ARMS | 27/24 | 9/8 | 0.4072 ± 0.1376 | 0.2283 ± 0.155 | 3.354 ± 0.06593 | 0.6192 |
| FC | MFMIHIFIS-197 | Sex + age | 3.50 | 2 | right | CTRL > ARMS | 27/24 | 5/4 | 0.1032 ± 0.1452 | −0.08055 ± 0.1137 | 4.232 ± 0.3204 | 0.5628 |
| FC | MFMIHIFIS-197 | Sex + age | 3.75 | 2 | right | CTRL > ARMS | 27/24 | 5/4 | 0.1032 ± 0.1452 | −0.08055 ± 0.1137 | 4.232 ± 0.3204 | 0.3216 |
| FC | MFMIHIFIS-197 | Sex + age | 4.00 | 1 | right | CTRL > ARMS | 27/24 | 4/3 | 0.1375 ± 0.1547 | −0.046 ± 0.128 | 4.377 ± 0.1658 | 0.2392 |
| FC | SFMIHIFIS-200 | Sex + age | 3.00 | 1 | right | CTRL > ARMS | 27/24 | 107/181 | 0.3199 ± 0.09473 | 0.1544 ± 0.0877 | 3.35 ± 0.3296 | 0.0988 |
| FC | SFMIHIFIS-200 | Sex + age | 3.25 | 1 | right | CTRL > ARMS | 27/24 | 77/91 | 0.3269 ± 0.08909 | 0.1523 ± 0.08574 | 3.573 ± 0.3206 | 0.0924 |
| FC | SFMIHIFIS-200 | Sex + age | 3.50 | 2 | right | CTRL > ARMS | 27/24 | 22/22 | 0.5063 ± 0.09474 | 0.3095 ± 0.09984 | 3.864 ± 0.3437 | 0.1464 |
| FC | SFMIHIFIS-200 | Sex + age | 3.75 | 3 | right | CTRL > ARMS | 27/24 | 10/9 | 0.5629 ± 0.1117 | 0.3538 ± 0.114 | 4.184 ± 0.3079 | 0.1472 |
| FC | SFMIHIFIS-200 | Sex + age | 4.00 | 1 | right | CTRL > ARMS | 27/24 | 6/5 | 0.5423 ± 0.1205 | 0.3135 ± 0.1322 | 4.395 ± 0.2421 | 0.1352 |
| **Structural connectivity (SC)** | | | | | | | | | | | | |
| SC | LFMIHIFIS-3 | Sex + age | 3.00 | 6 | right | CTRL > ARMS | 31/29 | 3/2 | 0.1567 ± 0.1197 | 0.07386 ± 0.08349 | 3.058 ± 0.03747 | 0.7544 |
| SC | LFMIHIFIS-3 | Sex + age | 3.25 | 1 | right | CTRL > ARMS | 31/29 | 2/1 | 0.2729 ± 0.2614 | 0.1019 ± 0.1252 | 3.355 | 1 |
| SC | LFMIHIFIS-3 | Sex + age | 3.50 | 0 | both | none above $\lvert t_0 \rvert$ | 31/29 | 0/0 | – | – | – | n/a |
| SC | LFMIHIFIS-3 | Sex + age | 3.75 | 0 | both | none above $\lvert t_0 \rvert$ | 31/29 | 0/0 | – | – | – | n/a |
| SC | LFMIHIFIS-3 | Sex + age | 4.00 | 0 | both | none above $\lvert t_0 \rvert$ | 31/29 | 0/0 | – | – | – | n/a |
| SC | MFMIHIFIS-197 | Sex + age | 3.00 | 1 | right | CTRL > ARMS | 30/29 | 2/1 | 0.004939 ± 0.007916 | 0.0009968 ± 0.002106 | 3.275 | 1 |
| SC | MFMIHIFIS-197 | Sex + age | 3.25 | 1 | right | CTRL > ARMS | 30/29 | 2/1 | 0.004939 ± 0.007916 | 0.0009968 ± 0.002106 | 3.275 | 1 |
| SC | MFMIHIFIS-197 | Sex + age | 3.50 | 1 | left | ARMS > CTRL | 30/29 | 2/1 | 0.06733 ± 0.1233 | 0.1027 ± 0.1092 | −3.552 | 0.798 |
| SC | MFMIHIFIS-197 | Sex + age | 3.75 | 0 | both | none above $\lvert t_0 \rvert$ | 30/29 | 0/0 | – | – | – | n/a |
| SC | MFMIHIFIS-197 | Sex + age | 4.00 | 0 | both | none above $\lvert t_0 \rvert$ | 30/29 | 0/0 | – | – | – | n/a |
| SC | SFMIHIFIS-200 | Sex + age | 3.00 | 4 | left | ARMS > CTRL | 31/28 | 4/3 | 0.9282 ± 0.4435 | 1.236 ± 0.5685 | −3.078 ± 0.08087 | 0.35 |
| SC | SFMIHIFIS-200 | Sex + age | 3.25 | 1 | left | ARMS > CTRL | 31/28 | 2/1 | 0.003297 ± 0.008116 | 0.005821 ± 0.01005 | −3.626 | 1 |
| SC | SFMIHIFIS-200 | Sex + age | 3.50 | 1 | left | ARMS > CTRL | 31/28 | 2/1 | 0.003297 ± 0.008116 | 0.005821 ± 0.01005 | −3.626 | 0.8688 |
| SC | SFMIHIFIS-200 | Sex + age | 3.75 | 0 | both | none above $\lvert t_0 \rvert$ | 31/28 | 0/0 | – | – | – | n/a |
| SC | SFMIHIFIS-200 | Sex + age | 4.00 | 0 | both | none above $\lvert t_0 \rvert$ | 31/28 | 0/0 | – | – | – | n/a |

### Structural context of the verified metabolic NBS component

Supplementary Tables S2 and S3 report the structural-context analysis of main-text Fig. 2 for the sex- and age-adjusted MC components of all three parcellations. The structural reference was the edge-wise mean structural connectome of the 30 healthy controls with matched MC data (29 for MFMIHIFIS-197). Hubs were defined within each hemisphere, and only intrahemispheric NBS edges entered the edge-class analysis (see main-text Methods).

Table S2: **Structural node strength of metabolic NBS nodes across parcellations and primary thresholds.** Sex- and age-adjusted MC NBS components (HR > HC). $p_{\mathrm{NBS}}$: two-direction-corrected component p-value (Table S1). Edges are split into left-intrahemispheric (LH), right-intrahemispheric (RH), and interhemispheric (inter). $n_{\mathrm{HC}}$: healthy controls with matched MC and SC data whose structural matrices were averaged to form the reference connectome. MW $p$ and $r_{\mathrm{rb}}$: two-sided Mann–Whitney test and rank-biserial correlation comparing structural strength of NBS and non-NBS nodes; $\rho$ and $p_{\mathrm{perm}}$: Spearman correlation between NBS node load and structural strength and its permutation p-value (5,000 permutations). Bold: $p < 0.05$, uncorrected across rows.

| Parcellation | $t_0$ | $p_{\mathrm{NBS}}$ | Nodes | Edges | LH | RH | Inter | $n_{\mathrm{HC}}$ | MW $p$ | $r_{\mathrm{rb}}$ | $\rho$ ($p_{\mathrm{perm}}$) |
|---|---|---|---|---|---|---|---|---|---|---|---|
| LFMIHIFIS-3 | 3.00 | 0.028 | 90 | 213 | 59 | 45 | 109 | 30 | < 0.001 | 0.30 | 0.24 (0.001) |
| | 3.25 | 0.026 | 68 | 120 | 31 | 23 | 66 | 30 | < 0.001 | 0.30 | 0.24 (0.002) |
| | 3.50 | 0.021 | 47 | 67 | 19 | 12 | 36 | 30 | 0.015 | 0.24 | 0.17 (0.017) |
| | 3.75 | 0.013 | 34 | 39 | 15 | 8 | 16 | 30 | 0.024 | 0.25 | 0.16 (0.036) |
| | 4.00 | 0.007 | 22 | 24 | 12 | 3 | 9 | 30 | 0.339 | 0.13 | 0.07 (0.356) |
| MFMIHIFIS-197 | 3.00 | 0.042 | 60 | 107 | 37 | 18 | 52 | 29 | 0.096 | 0.16 | 0.13 (0.097) |
| | 3.25 | 0.046 | 41 | 54 | 22 | 4 | 28 | 29 | 0.458 | 0.08 | 0.07 (0.389) |
| | 3.50 | 0.038 | 26 | 27 | 9 | 1 | 17 | 29 | 0.158 | 0.18 | 0.11 (0.166) |
| | 3.75 | 0.068 | 29 | 22 | 5 | 3 | 14 | 29 | 0.442 | 0.09 | 0.07 (0.383) |
| | 4.00 | 0.804 | 12 | 6 | 1 | 1 | 4 | 29 | 0.381 | 0.15 | 0.07 (0.397) |
| SFMIHIFIS-200 | 3.00 | 0.024 | 104 | 248 | 54 | 53 | 141 | 30 | < 0.001 | 0.43 | 0.34 (< 0.001) |
| | 3.25 | 0.025 | 79 | 136 | 25 | 30 | 81 | 30 | < 0.001 | 0.41 | 0.31 (< 0.001) |
| | 3.50 | 0.025 | 46 | 69 | 12 | 11 | 46 | 30 | 0.011 | 0.25 | 0.17 (0.019) |
| | 3.75 | 0.017 | 30 | 38 | 10 | 6 | 22 | 30 | 0.031 | 0.25 | 0.15 (0.030) |
| | 4.00 | 0.042 | 11 | 10 | 4 | 1 | 5 | 30 | 0.968 | 0.01 | 0.00 (0.992) |

Table S3: **Hub-node and edge-class enrichment of the metabolic NBS component at $t_0 = 3.25$ across parcellations.** Hubs were defined within each hemisphere as the top 2%, 5%, 10%, or 20% of control-average structural node strength. Hub nodes: NBS nodes that are hubs, compared with 5,000 random node sets of equal size from the same hemisphere. Edge classes: intrahemispheric NBS edges linking two hubs (hub–hub), one hub (feeder), or no hub (local), compared with 5,000 random intrahemispheric edge sets of equal size. obs: observed count; ER: enrichment ratio (observed / mean null); $p$: upper-tail empirical p-value. Bold: $p < 0.05$, uncorrected across thresholds, hemispheres, and parcellations.

| | | | | Hub nodes | | | Hub–hub | | Feeder | | | Local | | |
|---|---|---|---|---|---|---|---|---|---|---|---|---|---|---|
| Parcellation | Hemi. | Hubs | $n$ | obs | ER | $p$ | obs | $p$ | obs | ER | $p$ | obs | ER | $p$ |
| LFMIHIFIS-3 | LH | 2% | 2 | 0 | 0.00 | 1.000 | 0 | 1.000 | 0 | 0.00 | 1.000 | 31 | 1.04 | 0.263 |
| | | 5% | 5 | 2 | 1.30 | 0.489 | 0 | 1.000 | 8 | 2.53 | 0.011 | 23 | 0.83 | 0.997 |
| | | 10% | 10 | 3 | 0.97 | 0.653 | 0 | 1.000 | 12 | 2.01 | 0.009 | 19 | 0.77 | 0.995 |
| | | 20% | 19 | 8 | 1.36 | 0.173 | 0 | 1.000 | 19 | 1.88 | 0.001 | 12 | 0.61 | 0.998 |
| | RH | 2% | 2 | 0 | 0.00 | 1.000 | 0 | 1.000 | 0 | 0.00 | 1.000 | 23 | 1.05 | 0.369 |
| | | 5% | 5 | 0 | 0.00 | 1.000 | 0 | 1.000 | 0 | 0.00 | 1.000 | 23 | 1.12 | 0.078 |
| | | 10% | 10 | 3 | 1.45 | 0.333 | 0 | 1.000 | 5 | 1.11 | 0.476 | 18 | 0.99 | 0.662 |
| | | 20% | 19 | 6 | 1.53 | 0.160 | 1 | 0.612 | 9 | 1.18 | 0.328 | 13 | 0.90 | 0.809 |
| MFMIHIFIS-197 | LH | 2% | 2 | 1 | 1.70 | 0.507 | 0 | 1.000 | 3 | 2.69 | 0.092 | 19 | 0.91 | 0.978 |
| | | 5% | 4 | 2 | 1.70 | 0.348 | 0 | 1.000 | 7 | 3.23 | 0.005 | 15 | 0.76 | 0.999 |
| | | 10% | 8 | 3 | 1.27 | 0.440 | 0 | 1.000 | 8 | 1.95 | 0.043 | 14 | 0.79 | 0.982 |
| | | 20% | 16 | 6 | 1.27 | 0.308 | 1 | 0.591 | 10 | 1.38 | 0.153 | 11 | 0.79 | 0.928 |
| | RH | 2% | 2 | 0 | 0.00 | 1.000 | 0 | 1.000 | 0 | 0.00 | 1.000 | 4 | 1.06 | 0.798 |
| | | 5% | 4 | 1 | 3.55 | 0.266 | 0 | 1.000 | 1 | 2.32 | 0.369 | 3 | 0.84 | 0.939 |
| | | 10% | 8 | 1 | 1.77 | 0.446 | 0 | 1.000 | 1 | 1.23 | 0.590 | 3 | 0.95 | 0.803 |
| | | 20% | 15 | 1 | 0.95 | 0.710 | 0 | 1.000 | 1 | 0.74 | 0.806 | 3 | 1.21 | 0.517 |
| SFMIHIFIS-200 | LH | 2% | 2 | 0 | 0.00 | 1.000 | 0 | 1.000 | 0 | 0.00 | 1.000 | 25 | 1.04 | 0.360 |
| | | 5% | 5 | 0 | 0.00 | 1.000 | 0 | 1.000 | 0 | 0.00 | 1.000 | 25 | 1.11 | 0.072 |
| | | 10% | 10 | 2 | 0.90 | 0.706 | 0 | 1.000 | 8 | 1.74 | 0.071 | 17 | 0.84 | 0.964 |
| | | 20% | 20 | 5 | 1.12 | 0.474 | 1 | 0.634 | 9 | 1.11 | 0.419 | 15 | 0.94 | 0.730 |
| | RH | 2% | 2 | 2 | 3.03 | 0.101 | 0 | 1.000 | 4 | 3.37 | 0.027 | 26 | 0.90 | 0.994 |
| | | 5% | 5 | 4 | 2.42 | 0.038 | 0 | 1.000 | 7 | 2.43 | 0.024 | 23 | 0.85 | 0.991 |
| | | 10% | 10 | 6 | 1.82 | 0.060 | 1 | 0.237 | 7 | 1.28 | 0.308 | 22 | 0.91 | 0.898 |
| | | 20% | 20 | 9 | 1.36 | 0.149 | 1 | 0.695 | 12 | 1.24 | 0.239 | 17 | 0.89 | 0.843 |

### Metabolite contrasts underlying the MC modes

To identify which metabolites drive the first two MC modes, $z$-scored control-average maps of tCr, Glx, Cho, tNAA, and Ins were combined pairwise into common maps, $(z_A + z_B)/2$, and contrast maps, $(z_A - z_B)/2$, and each map was correlated with $g_1$ and $g_2$ across parcels (see main-text Methods). $g_1$ was dominated by contrasts of Cho against tNAA and Ins, and $g_2$ by contrasts of Glx against Cho and tNAA. Common-map correlations were weak for both modes (all $|r| \leq 0.43$), indicating that the modes reflect relative differences between metabolites rather than overall metabolite abundance.

Table S4: **Correlations of metabolite-pair contrast and common maps with MC modes $g_1$ and $g_2$** (LFMIHIFIS-3, healthy controls). Contrast: $(z_A - z_B)/2$; common: $(z_A + z_B)/2$.

| | $g_1$ | | $g_2$ | |
|---|---|---|---|---|
| Metabolite pair ($A$ vs $B$) | contrast | common | contrast | common |
| Cho vs tNAA | −0.87 | −0.02 | 0.09 | 0.27 |
| Cho vs Ins | −0.85 | −0.09 | 0.32 | 0.13 |
| tCr vs Cho | 0.77 | −0.14 | −0.37 | 0.11 |
| Glx vs Cho | 0.69 | −0.17 | −0.57 | −0.02 |
| Glx vs tNAA | −0.33 | 0.36 | −0.62 | −0.07 |
| Glx vs Ins | −0.30 | 0.29 | −0.51 | −0.18 |
| tCr vs tNAA | −0.36 | 0.38 | −0.48 | 0.04 |
| tCr vs Glx | 0.12 | 0.24 | 0.46 | −0.20 |
| tNAA vs Ins | 0.20 | 0.43 | 0.41 | 0.07 |
| tCr vs Ins | −0.28 | 0.31 | −0.13 | −0.07 |

Table S5: **Per-metabolite participation in the contrast structure of MC modes $g_1$ and $g_2$** (LFMIHIFIS-3, healthy controls). For each metabolite, the signed column averages its signed contrast correlation $(z_A - z_B)/2$ over the four pairs in which it appears (sign flipped when the metabolite is the second-named partner $B$), giving its net relative direction along the mode; the magnitude column averages $|r|$ over the same four pairs, giving its overall contribution irrespective of direction. Metabolites are ranked by magnitude within each mode.

| $g_1$ | | | $g_2$ | | |
|---|---|---|---|---|---|
| Metabolite | signed | magnitude | Metabolite | signed | magnitude |
| Cho | −0.797 | 0.797 | Glx | −0.539 | 0.539 |
| tNAA | 0.441 | 0.441 | tNAA | 0.353 | 0.398 |
| Ins | 0.308 | 0.407 | tCr | −0.130 | 0.360 |
| tCr | 0.062 | 0.385 | Ins | −0.021 | 0.342 |
| Glx | −0.014 | 0.361 | Cho | 0.337 | 0.337 |

On $g_1$, Cho dominates (signed $r = -0.80$), decreasing relative to the other four metabolites toward higher mode values, while tNAA and Ins increase; tCr and Glx contribute comparatively little. On $g_2$, Glx is the strongest metabolite, decreasing toward higher mode values (signed $r = -0.54$), while Cho and tNAA increase; tCr and Ins contribute weakly. Cho therefore anchors the decreasing pole of $g_1$, opposed to tNAA and Ins, but the increasing pole of $g_2$, together with tNAA and opposed to Glx.

### Mathematical Note S1: A network-level variational argument for spectral MC modes

This note provides an idealized mathematical motivation for the curved two-dimensional geometry formed by the first two metabolic connectivity (MC) modes. It is not a proof that an empirical MC embedding must be triangular. Rather, it shows that, under explicit locality and smooth-domain assumptions, a network balancing grey-matter-constrained local coupling against global edge diversity leads to smooth spectral modes. The first two modes then form a parabolic relation when they behave as successive harmonics of a shared intrinsic coordinate; anatomical heterogeneity and branching can broaden this curve into the approximately triangular point cloud observed empirically.

#### Local geometric coupling and global edge diversity

Let $G = (V, E)$ be a graph with nodes $V = \{1, \ldots, N\}$ representing brain parcels and edges $E$ representing admissible interactions along grey-matter-continuous geometry. We consider symmetric non-negative network weights $W_{ij} = W_{ji} \geq 0$ for $(i, j) \in E$, normalized such that

$$\sum_{(i,j)\in E} W_{ij} = 1.$$

Each admissible edge is assigned a non-negative geometric cost $c_{ij}$. For example, $c_{ij}$ may be the squared Euclidean, geodesic, or tissue-constrained distance between parcels, with non-admissible pairs excluded from $E$. The geometric cost and edge-weight entropy are

$$E_{\mathrm{geo}}[W] = \sum_{(i,j)\in E} W_{ij} c_{ij}, \qquad H[W] = -\sum_{(i,j)\in E} W_{ij} \log W_{ij}.$$

We model the network as minimizing geometric cost while retaining globally diverse edge weights:

$$W^\star = \arg\min_W \left[ E_{\mathrm{geo}}[W] - \tau H[W] \right],$$

subject to symmetry, non-negativity, and normalization. The parameter $\tau > 0$ controls the balance: smaller $\tau$ gives greater preference to low-cost local interactions, whereas larger $\tau$ permits a broader distribution of weights over admissible edges.

Introducing a Lagrange multiplier $\alpha$ for normalization gives

$$\mathcal{G}[W] = \sum_{(i,j)\in E} W_{ij} c_{ij} + \tau \sum_{(i,j)\in E} W_{ij} \log W_{ij} + \alpha \left( \sum_{(i,j)\in E} W_{ij} - 1 \right).$$

Stationarity with respect to $W_{ij}$ requires

$$\frac{\partial \mathcal{G}}{\partial W_{ij}} = c_{ij} + \tau(\log W_{ij} + 1) + \alpha = 0,$$

and therefore

$$W_{ij} = C \exp\left( -\frac{c_{ij}}{\tau} \right).$$

Applying the normalization constraint yields

$$\boxed{W^\star_{ij} = \frac{1}{Z} \exp\left( -\frac{c_{ij}}{\tau} \right)}, \qquad Z = \sum_{(i,j)\in E} \exp\left( -\frac{c_{ij}}{\tau} \right).$$

The optimized network is thus a distance-decaying Gibbs kernel: it favors local interactions defined by grey-matter-continuous geometry while entropy prevents all weight from collapsing onto only the minimum-cost edge.

#### From graph smoothness to the Neumann eigenproblem

Let

$$L^\star = D^\star - W^\star, \qquad D^\star_{ii} = \sum_j W^\star_{ij}$$

be the graph Laplacian of the optimized network. For a scalar field $\phi : V \to \mathbb{R}$, its graph Dirichlet energy is

$$Q_{W^\star}[\phi] = \frac{1}{2}\sum_{i,j} W^\star_{ij}(\phi_i - \phi_j)^2.$$

The nonconstant graph modes minimize the Rayleigh quotient

$$\mathcal{R}_{W^\star}[\phi] = \frac{Q_{W^\star}[\phi]}{\sum_i(\phi_i - \bar{\phi})^2}, \qquad \bar{\phi} = \frac{1}{N}\sum_i \phi_i,$$

subject to orthogonality to the constant mode and, for subsequent modes, to the preceding eigenvectors. Equivalently, they solve

$$L^\star g_k = \lambda_k g_k.$$

Now assume that parcels densely sample a smooth domain $\Omega \subset \mathbb{R}^3$ and that the effective distance-decaying kernel is local and symmetric. The first-order term in the local expansion of the nonlocal graph energy then vanishes by symmetry, while the leading nonzero term is proportional to continuum Dirichlet energy:

$$Q_{W^\star}[\phi] \approx C_\tau \int_\Omega \|\nabla\phi(\mathbf{r})\|^2 \, d\mathbf{r},$$

where $C_\tau > 0$ depends on the kernel scale. After centering $\phi$, the corresponding continuum quotient is

$$\mathcal{R}[\phi] = \frac{\int_\Omega \|\nabla\phi(\mathbf{r})\|^2 \, d\mathbf{r}}{\int_\Omega \phi(\mathbf{r})^2 \, d\mathbf{r}}.$$

At a stationary point, an arbitrary perturbation $\eta$ satisfies

$$\int_\Omega \nabla\phi \cdot \nabla\eta \, d\mathbf{r} = \widetilde{\lambda} \int_\Omega \phi\eta \, d\mathbf{r}.$$

Integration by parts gives

$$\int_\Omega \nabla\phi \cdot \nabla\eta \, d\mathbf{r} = -\int_\Omega (\Delta\phi)\eta \, d\mathbf{r} + \int_{\partial\Omega} \eta \, \nabla\phi \cdot \mathbf{n} \, dS.$$

Because field values are not fixed at the boundary, the boundary perturbation is arbitrary and the natural boundary condition is

$$\nabla\phi \cdot \mathbf{n} = 0 \qquad \text{on } \partial\Omega.$$

The interior condition is consequently

$$-\Delta\phi = \widetilde{\lambda}\phi \qquad \text{in } \Omega.$$

Thus, after absorbing $C_\tau$ into the eigenvalues, minimizing local smoothness per unit global variance yields the Neumann eigenproblem

$$\boxed{-\Delta g_k = \widetilde{\lambda}_k g_k \quad \text{in } \Omega, \qquad \nabla g_k \cdot \mathbf{n} = 0 \quad \text{on } \partial\Omega}.$$

Its zero-eigenvalue solution is constant. After centering, $g_1$ is the smoothest nonconstant spatial contrast, and subsequent modes are the smoothest contrasts orthogonal to the preceding modes.

#### First and second harmonics and the triangular approximation

Consider first a one-dimensional intrinsic coordinate $s \in [0, L]$. The Neumann eigenfunctions are

$$\psi_0(s) = \frac{1}{\sqrt{L}}, \qquad \psi_{k\geq 1}(s) = \sqrt{\frac{2}{L}} \cos\left(\frac{k\pi s}{L}\right), \qquad \lambda_k = \left(\frac{k\pi}{L}\right)^2.$$

Writing $\theta = \pi s/L$, the first two centered and normalized nonconstant modes behave as

$$\widetilde{g}_1 \approx \cos\theta, \qquad \widetilde{g}_2 \approx \cos 2\theta.$$

Using $\cos 2\theta = 2\cos^2\theta - 1$ eliminates the latent coordinate and gives

$$\boxed{\widetilde{g}_2 \approx 2\widetilde{g}_1^2 - 1}.$$

The ideal two-mode embedding is therefore parabolic. The two extrema of $\widetilde{g}_1$ define opposite endpoint states, while the extremum of $\widetilde{g}_2$ defines the bend of the curve.

In a folded three-dimensional domain, $g_1$ does not impose a unique global ordering of all parcels. Instead, it defines isolevel surfaces

$$\Sigma_c = \{\mathbf{r} \in \Omega : g_1(\mathbf{r}) = c\}$$

and gradient-flow trajectories $\gamma_{\mathbf{r}_0}(s)$ satisfying

$$\frac{d\gamma_{\mathbf{r}_0}}{ds} = \frac{\nabla g_1(\gamma_{\mathbf{r}_0}(s))}{\|\nabla g_1(\gamma_{\mathbf{r}_0}(s))\|}.$$

Along each trajectory for which $\nabla g_1 \neq 0$,

$$\frac{d}{ds} g_1(\gamma_{\mathbf{r}_0}(s)) = \|\nabla g_1(\gamma_{\mathbf{r}_0}(s))\| \geq 0.$$

Monotonicity is therefore recovered pathwise rather than as one global parcel ordering. If the first two modes are approximately successive harmonics of a shared coordinate along these trajectories, their point cloud retains the parabolic tendency derived above. Finite parcellation, anatomical heterogeneity, multiple branches, and deviations from a single shared coordinate broaden this curve. The maximal-area triangle fitted to the empirical convex hull is consequently used as a piecewise-linear summary of this broadened geometry, not as a boundary condition or a theoretical requirement of diffusion embedding.
